# Genomic compartmentalization of pervasive sex-biased gene expression in the vine mealybug *Planococcus ficus*

**DOI:** 10.64898/2026.07.01.735863

**Authors:** Dario Cantu, Rosa Figueroa-Balderas, Mark Sisterson, Andrea Minio, Noé Cochetel, Rachel Naegele, Lindsey Burbank

**Affiliations:** Department of Viticulture and Enology, University of California Davis, Davis CA 95618, USA; Genome Center, University of California Davis, Davis CA 95618, USA; USDA Agricultural Research Service, San Joaquin Valley Agricultural Sciences Center, Parlier, CA, USA; USDA Agricultural Research Service, Sugarbeet and Bean Research Unit, East Lansing, MI 48824, U.S.A

**Keywords:** paternal genome elimination, sex-biased gene expression, sexual dimorphism, *Planococcus ficus*, genome assembly, salivary effectors, vine mealybug

## Abstract

The vine mealybug, *Planococcus ficus*, is a globally invasive pest of grapevine and a vector of leafroll viruses. Like other mealybugs, it reproduces through paternal genome elimination, a sex-determination system that operates without sex chromosomes and is associated with extreme sexual dimorphism. To characterize genome organization and sex-biased expression in this species, we generated a long-read reference genome spanning 369 Mb with 23,489 annotated genes and macrosynteny conserved with the citrus mealybug, *Planococcus citri*. Resequencing of four California field individuals yielded a first whole-genome estimate of nucleotide diversity and 132 microsatellite markers for population monitoring. Among 2,129 candidate secreted proteins, a conserved core is shared with *P. citri*, but each species carries a distinct set of lineage-specific effectors. Comparing adult male and female transcriptomes, we found sex-biased expression to be pervasive and skewed toward females: 41% of tested genes differed between the sexes, with female-biased genes both more numerous and showing larger fold changes. These female-biased genes were not randomly distributed but concentrated in discrete blocks of coordinately expressed, tandemly duplicated gene families, a pattern not previously described in a mealybug. Male- and female-biased secreted proteins also differed in origin, with male-biased proteins drawn from a conserved repertoire shared with *P. citri* and female-biased proteins spanning a more lineage-specific pool. Together, these results reveal a female-skewed, spatially clustered architecture of sex-biased expression in a mealybug that lacks sex chromosomes, and provide genomic resources for managing an invasive vineyard pest.

## Introduction

The vine mealybug, *Planococcus ficus* (Signoret) (Hemiptera: Pseudococcidae), is a major pest of grapevine worldwide. Native to the Mediterranean basin, the Middle East, and parts of northern Africa, it has invaded California, Mexico, South America, and southern Africa over the past few decades (Daane et al. 2018; Cocco et al. 2021). Damage is both direct and indirect: heavy infestations reduce vine vigor and contaminate clusters with honeydew and sooty mold, while transmission of grapevine leafroll-associated virus 3 (GLRaV-3) and other ampeloviruses causes the greatest economic losses (Tsai et al. 2008; Tsai et al. 2010). Management relies heavily on a narrow set of insecticide chemistries, raising concerns about resistance, non-target effects, and sustainability (Mansour et al. 2018; Daane et al. 2020). *P. ficus* overwinters under the bark and moves into the canopy as the season progresses, making timing and coverage of contact treatments difficult (Sisterson and Uchima 2024). Developing alternative control strategies, including RNAi, biocontrol, and resistant rootstocks, requires genomic resources that are still limited for this species.

*P. ficus* belongs to the Pseudococcidae, a family of roughly 2,000 unarmored scale insects within the superfamily Coccoidea (Hemiptera). Relationships among the major lineages of Coccoidea have been substantially clarified by recent molecular and genomic phylogenies (Choi and Lee 2022), but divergence times within Pseudococcidae remain less well characterized, with few genome-based estimates for the major pest genera (**Fig. 1**). The genus *Planococcus* includes several agricultural pests in addition to *P. ficus*, most notably *P. citri* (citrus mealybug) and *P. minor* (passionvine mealybug). *P. ficus* overlaps with *P. citri* in host range, and the three species are difficult to separate by morphology alone, often requiring molecular markers for reliable identification (Daane et al. 2011). Genome assemblies are now available for more than a dozen Coccoidea species, including chromosome-level assemblies for several Pseudococcidae and Coccidae (M. Li et al. 2020; Ross et al. 2024), providing a comparative framework that until recently was limited to a handful of draft genomes. Population genetic studies indicate that invasive *P. ficus* populations, including those in California, carry reduced diversity consistent with founder effects (Daane et al. 2018).

**Figure 1.**
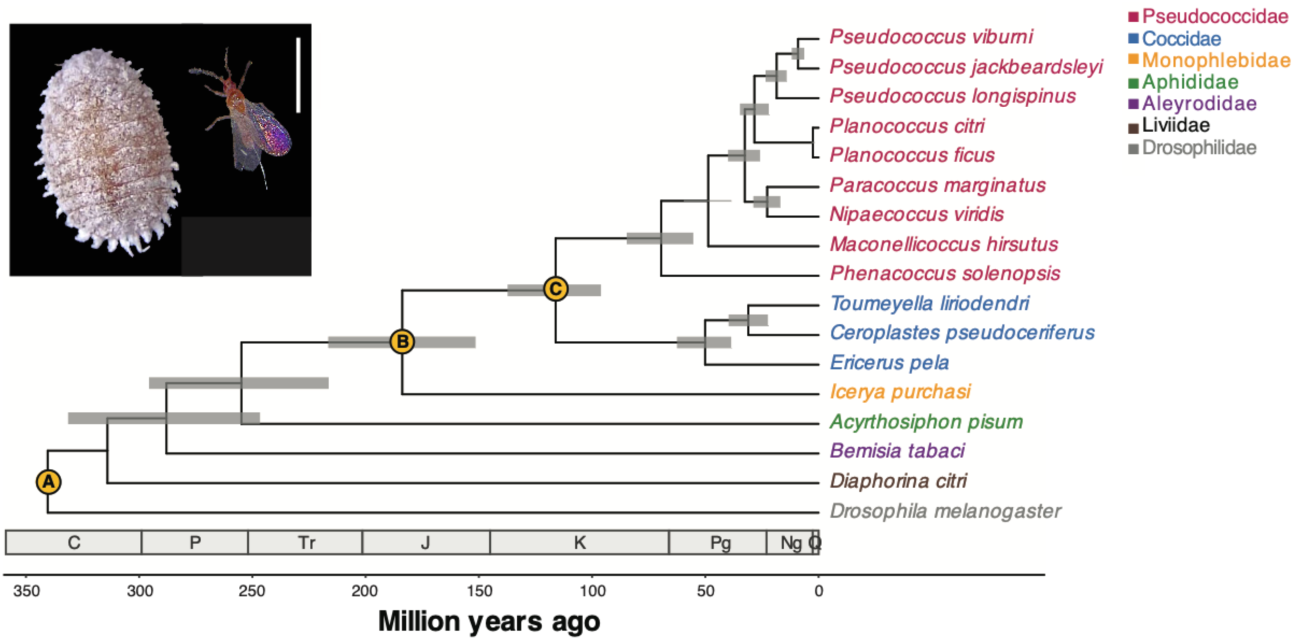
Time-calibrated phylogeny of *Planococcus ficus* (male and female in photo insets, left and right, respectively; photos by Zachary Dashner) and 16 related Hemiptera, rooted with *Drosophila melanogaster*. Maximum-likelihood topology inferred with IQ-TREE under LG+R4 from a concatenated amino-acid supermatrix; all internal nodes received 100% SH-aLRT and 98–100% UFBoot support. Divergence times were estimated with MCMCTree (PAML 4.9j) under an independent-rates relaxed clock, calibrated at three fossil-anchored nodes (gold circles): node A, Diptera–Hemiptera split, soft bounds 300–365 Ma (Misof et al. 2014); node B, Coccoidea crown, minimum 130 Ma from *Hodgsonicoccus patefactus* in Lebanese amber (Vea and Grimaldi 2015); node C, Neococcoidea crown, minimum 99 Ma from *Kozarius perpetuus* and *Rosahendersonia prisca* in Burmese amber (Shi et al. 2012; Vea and Grimaldi 2015). Minimum-age constraints were applied as soft lower bounds with 2.5% tail probability. Grey bars: 95% HPD intervals. Tip labels colored by family. Geological periods: C, Carboniferous; P, Permian; Tr, Triassic; J, Jurassic; K, Cretaceous; Pg, Paleogene; Ng, Neogene; Q, Quaternary.

Mealybugs exhibit paternal genome elimination (PGE), a genetic system in which both sexes are diploid at fertilization but the paternally inherited chromosome set is heterochromatinized and largely transcriptionally silenced in males during early embryogenesis (Kol-Maimon et al. 2014; Bain et al. 2021; de la Filia et al. 2021; Herbette and Ross 2023). Males are therefore often described as functionally haploid, although somatic paternal silencing is in practice partial (de la Filia et al. 2021). During a highly modified, inverted meiosis the paternal chromosomes are partitioned into degenerating spermatids, so males transmit almost exclusively their maternally derived genome, with rare leakage of single paternal chromosomes (Bongiorni et al. 2004; de la Filia et al. 2019). PGE is widespread in arthropods, where it has evolved independently at least six times and occurs in over 10,000 species (Gardner and Ross 2014; de la Filia et al. 2015; Herbette and Ross 2023). This reproductive strategy has implications for the evolution of sex-biased gene expression, dosage compensation, and pesticide resistance (Brun et al. 1995; Carrière 2003). Sex-specific transcriptomes of *P. ficus* are expected to reveal how extensively expression differs between the two sexes and how that variation is distributed across the genome, providing a framework for interpreting sex-biased expression in a species whose sex arises under PGE rather than through sex chromosomes (Herbette and Ross 2023).

Like other phloem-feeding hemipterans, *P. ficus* injects saliva into host tissues during feeding, delivering a complex mixture of proteins and small molecules that modulate plant defenses, alter phloem sieve element function, and facilitate sustained ingestion (Will et al. 2013). In aphids, whiteflies, and psyllids, these salivary effectors include enzymes that degrade or modify plant cell walls, such as pectinases and cellulases (McAllan and Adams 1961; Ma et al. 1990; Sharma et al. 2014; Silva-Sanzana et al. 2020), calcium-binding proteins that prevent sieve tube occlusion (Will et al. 2007), and small secreted proteins that suppress jasmonate- and salicylate-mediated immunity (Elzinga and Jander 2013); the mealybug *Phenacoccus solenopsis* likewise suppresses host defenses by manipulating jasmonate-salicylate crosstalk (Zhang et al. 2015). The effector repertoire of mealybugs nonetheless remains poorly characterized compared with that of aphids (Carolan et al. 2011), despite the economic importance of the group and its role as a virus vector. Cataloging predicted secreted proteins in the *P. ficus* genome and comparing them to those of *P. citri* and other Coccoidea will identify conserved core effectors, lineage-specific expansions, and candidates that may underlie host range differences or interactions with grapevine leafroll viruses during transmission (Tsai et al. 2008; Almeida et al. 2013).

Here we present a long-read reference genome for *P. ficus*, annotate its candidate secreted-protein repertoire in comparison with *P. citri*, benchmark genetic diversity in California populations and develop cross-validated SSR markers, and map sex-biased gene expression genome-wide. We show that sex-biased expression is pervasive and female-skewed, that female-biased genes are organized into discrete genomic blocks of coordinately co-expressed tandem paralogs, and that the male- and female-biased secretomes differ in evolutionary character. Together these resources and analyses establish a genomic foundation for *P. ficus* and place its sex-biased expression in the context of paternal genome elimination.

## Materials and Methods

### Biological material

Vine mealybugs used for reference genome sequencing (*P. ficus* VMBCA21f) and corresponding transcriptome sequencing were from a laboratory colony reared on squash for several years. Individual male and female insects for sequencing were grown separately as described below.Vine mealybugs sequenced for evaluation of genetic diversity (*P. ficus* MB21-2, MB21-3, MB21-P, and MB21-Q) were collected from four vineyards in the San Joaquin Valley of California (1 location near Parlier, CA, 2 locations near Jasmin, CA, and 1 location near Saco, CA) on August 12, 2021, and used to establish independent colonies. For field collected insects,vine mealybug infested grape clusters were transported from the field to the laboratory in a cooler and placed in 60 x 43 x 15 cm trays lined with paper towels. Squash (butternut or acorn) was placed in the tray with the infested grape clusters. As the grape clusters dried down, the mealybugs moved off the grape clusters and onto the squash. Once enough mealybugs had moved onto the squash, the mealybug infested squash were placed in 40 x 50 x 36 cm storage totes lined with vermiculate to absorb moisture as described by Sisterson and Uchima (2024). Each tote was placed on a mat encircled with Stikem special (Seabright laboratories, Emeryville, CA) to prevent movement of crawlers. Colonies were held at room temperature (∼23°C) and ambient light. To generate virgins for sequencing, grape seedlings (∼15-20 cm tall, open pollinated ‘Chardonnay’) were placed on top of colony cages where crawlers actively roamed. Once seedlings were infested with ∼100 crawlers, the grape seedlings were moved to a plant growth chamber and held for 9-12 days at 27°C with 14:10 L:D cycle. After 9-12 days, the grape seedlings were cut into pieces and placed into 11 cm radius deli containers (Amerifoods Trading Company, Los Angeles, CA). As the plant material in the deli container dried, the mealybugs released their mouth parts from the plant pieces and began actively roaming the container. Individual mealybugs were transferred to ∼ 5 cm tall grape seedlings using a fine paint brush and sealed in the cages described by Sisterson and Uchima (2024). Cages with individual mealybugs were held for an additional 2 weeks at 27°C and 14:10 L:D cycle for mealybugs to reach maturity. Because males typically pupate around 14 days under these conditions Sisterson and Uchima (2024), insects were separated and held individually prior to reaching reproductive maturity.

### DNA library preparation and sequencing

High-Molecular Weight genomic DNA from single female insect was isolated using the Quick DNA HMW MagBead kit (Zymo Research, Irvine, CA, USA). DNA purity, quantity and integrity were evaluated with a Nanodrop 2000 spectrophotometer (Thermo Scientific, Hanover Park, IL, USA), with the DNA High Sensitivity kit on a Qubit 2.0 Fluorometer (Life Technologies, Carlsbad, CA, USA), and by FEMTO Pulse System (Agilent, Santa Clara, CA, USA), respectively. HiFi libraries were prepared following the PacBio Ultra-Low DNA Input Library Preparation using the SMRTbell Express Template Prep Kit 2.0 protocol (PN 101-987-800, Version 01, Pacific Biosciences, Menlo Park, CA, USA). Briefly, 10 ng of HMW gDNA was sheared to a modal size of 10-15 Kb using Diagenode’s Megaruptor (Diagenode LLC, Denville, NJ, USA) and subjected to DNA damage repair and end repair/A-tailing. PCR adapters were then ligated to the repaired fragments, followed by 15 cycles of PCR amplification to generate sufficient material for library construction. The amplified DNA was pooled, subjected to a second round of DNA damage repair and end repair/A-tailing, and converted into SMRTbell libraries through ligation of SMRTbell adapters. Libraries were purified and size-selected using the BluePippin instrument (Sage Science, Beverly, MA, USA) with a size selection range of 8-17 Kb. The size-selected library was cleaned using 1x ProNex beads (Promega, Madison, WI, USA), and its concentration and final size distribution were evaluated using a Qubit HS Assay Kit (Thermo Scientific, Hanover Park, IL, USA) and the Femto Pulse System (Agilent, Santa Clara, CA, USA), respectively. The final library was sequenced on a PacBio Sequel IIe platform (DNA Technology Core Facility, University of California, Davis). For short read DNA sequencing, 200 ng of HMW gDNA was used for library preparation using the Kapa LTP library prep kit (Kapa Biosystems, Wilmington, MA, USA). The DNA library was evaluated for quantity and quality with the High Sensitive chip on a Bioanalyzer 2100 (Agilent Technologies, Santa Clara, CA, USA) and sequenced in paired-end 150 bp reads on an Illumina HiSeq X Ten instrument (IDseq, CA, USA).

### Reference genome sequencing and assembly

*De novo* assembly was performed with FALCON and FALCON-Unzip from falcon-kit 1.8.1 (Chin et al. 2016). In the read-correction stage, raw subreads were processed with a seed length cutoff of 100 bp. Pairwise overlaps were detected with daligner (-e0.7 -l100 -k16 -h64 -w7 -s100), tandem and interspersed repeats were masked with HPCTANmask and HPCREPmask (-k18 -h480 -w8 -e.8 -s100; REPmask_code 1,15;2,10;4,10), and error-corrected reads were generated with falcon_sense requiring at least 70% identity and 3x coverage (--min_idt 0.70 --min_cov 3 --max_n_read 400). In the assembly stage, error-corrected reads of at least 7 kb were used as input. Overlaps among corrected reads were computed with daligner (-t60 -k18 -h256 -e.8 -l100 -s100) and filtered with a highly heterozygous setting (--max_diff 100 --max_cov 400 --min_cov 4) to retain divergent allelic contigs, chosen given the heterozygosity range (0.18-0.30%) estimated from k-mer analysis. FALCON-Unzip partitioned the assembly into primary contigs (Ps) and haplotigs (Hc). Both haplotype sets were polished with Arrow using gcpp (GenomicConsensus, PacBio). Primary (Ps) contigs were scaffolded with SSPACE-LongRead (Boetzer and Pirovano 2014) using the PacBio subreads (BLASR alignment), with a minimum alignment identity of 70%, a minimum of 10 links, a link ratio of 0.3, and a minimum overlap of 10; haplotigs (Hc) were retained as unscaffolded contigs. The combined Ps + Hc assembly was carried forward for decontamination, annotation, and downstream analyses. Each scaffold was compared by nucleotide BLAST (blastn; NCBI BLAST+ 2.7.1) against a custom local database of reference genomes spanning bacteria, archaea, fungi, plants, and metazoans, and the best hit per scaffold was retained and classified by source organism. Scaffolds whose best hits were to non-arthropod genomes with high query coverage and identity were flagged as contamination. Flagged scaffolds, together with their associated gene, transcript, and repeat annotations, were removed prior to all downstream analyses.

Genome size, heterozygosity, and repeat content were estimated from Illumina short reads of four California *P. ficus* individuals. K-mers (k = 21) were counted with Jellyfish v.2.2.7 (Marçais and Kingsford 2011) and modeled with GenomeScope2 (v2.0; Ranallo-Benavidez et al. 2020) under a diploid model. Assembly statistics were computed with seqkit and samtools. Completeness was assessed with BUSCO v.6.0.0 (Manni et al. 2021) against the hemiptera_odb12 lineage, in genome mode (miniprot-based prediction) and protein mode (longest isoform per locus). The *P. ficus* assembly was aligned to the chromosome-level *P. citri* reference (GCF_950023065.1) with minimap2 v.2.28 (Li 2021) using the asm20 preset and --secondary=no. Alignments shorter than 2 kb or on unplaced *P. citri* scaffolds were discarded. Each *P. ficus* scaffold was assigned an orientation from its dominant alignment strand to *P. citri*, and collinear alignments within 1 Mb of one another on both query and target were chained into syntenic blocks. Gene-level synteny was independently assessed with MCScanX (Wang et al. 2012) from DIAMOND v.2.2.0 blastp alignments (e-value 1e-10; Buchfink et al. 2021) between the longest protein isoforms of the two species, using default MCScanX parameters (minimum 5 collinear genes per block).

### Gene prediction and functional annotation

Protein-coding genes were predicted with the EVidenceModeler-based pipeline of the Cantu lab (https://github.com/andreaminio/AnnotationPipeline-EVM_based-DClab; Cochetel et al. 2021) over the combined Ps + Hc assembly. *P. ficus* RNA-seq libraries described below were assembled *de novo* and genome-guided with Trinity v.2.6.5 (Haas et al. 2013) and, after HISAT2 v.2.0.5 alignment (Kim et al. 2015), with StringTie v.1.3.4d (Pertea et al. 2015); coding regions were extracted with TransDecoder v.3.0.1 (TransDecoder). Transcripts were aligned to the assembly with GMAP v.2015-09-29 (Wu and Watanabe 2005), BLAT v.36×2 (Kent 2002), and Magic-BLAST v.1.4.0 (Boratyn et al. 2019), and reconciled into alignment assemblies with PASA v.2.3.3 (Haas et al. 2003). Protein evidence was aligned with exonerate v.2.2.0 (Slater and Birney 2005). A training set derived from the PASA assemblies was used to train AUGUSTUS v.3.0.3 (Stanke et al. 2006), GeneMark.hmm v.3.47 (Lomsadze et al. 2005), and SNAP v.2006-07-28 (Korf 2004). Gene models were predicted with these *ab initio* predictors, a BUSCO-trained AUGUSTUS model (BUSCO v.3.0.2, insecta and eukaryota; Manni et al. 2021), and PASA/TransDecoder-derived models. All prediction tracks, transcript alignments and exonerate protein alignments were integrated with EVidenceModeler v.1.1.1 (Haas et al. 2008), weighting PASA assemblies most heavily (weight 20) and PASA/TransDecoder models (25), followed by *ab initio* predictions, transcript alignments, and protein alignments. Consensus models were polished and updated for UTRs and alternative isoforms with PASA, then filtered to remove transposable-element and low-confidence models using DIAMOND/BLAST against NCBI RefSeq invertebrate (Camacho et al. 2009). Functional annotation was assigned with Blast2GO v.4.1.9 (Götz et al. 2008) from DIAMOND/BLAST homology against NCBI RefSeq and InterProScan v.5.28-67.0 (Jones et al. 2014) domain assignments.

### Effector prediction

Signal peptides in the longest annotated isoform per gene of *P. ficus* and *P. citri* (NCBI RefSeq GCF_950023065.1; Ross et al. 2024) were predicted with SignalP 6.0 in eukaryotic fast mode (Teufel et al. 2022). Proteins with an SP probability ≥ 0.7 and a mature peptide length of 30-500 amino acids were retained as candidate effectors. Mature peptide length and cysteine content were computed directly from the SignalP-trimmed sequences. The two filtered secretomes were combined and clustered jointly. All-versus-all sequence similarity was computed with DIAMOND blastp v2.2.0 (E ≤ 1 x 10⁻⁵; Buchfink et al. 2021), and the resulting hits were clustered with MCL v22-282 (Van Dongen 2008) at inflation I = 2.0. Clustering yielded 1,596 groups spanning all 4,312 candidate effectors, including four proteins that had no significant similarity to any other sequence and were retained as single-member clusters. Clusters were categorized as shared 1:1, shared with expansion in one or both lineages, or lineage-specific (singleton or family expansion without a cross-species homolog). Protein-domain content of the candidate effectors was annotated with InterProScan v5.70-102.0 (Jones et al. 2014).

### Variant calling and preliminary genetic diversity

Whole-genome resequencing reads from four field-collected populations of *P. ficus* (MB21-2, MB21-3, MB21-P, and MB21-Q) were aligned to the *P. ficus* VMBCA21f primary assembly with BWA-MEM v0.7.10 (Li and Durbin 2009; Li 2013) and processed through duplicate marking. Per-sample variants were called with GATK HaplotypeCaller (v4.6.2.0; McKenna et al. 2010) in gVCF mode at diploid ploidy, then joint-genotyped with CombineGVCFs and GenotypeGVCFs. SNPs and indels were split and hard-filtered using the GATK best-practice thresholds for small cohorts (used in place of VQSR) (SNPs: QD<2, FS>60, MQ<40, MQRankSum<-12.5, ReadPosRankSum<-8, SOR>3; indels: QD<2, FS>200, ReadPosRankSum<-20, SOR>10). To enable callable-site denominators for nucleotide diversity estimation, GenotypeGVCFs was rerun with --include-non-variant-sites to produce an all-sites VCF; variant records were split, hard-filtered, and concatenated back with the invariant records using bcftools v1.23.1 (Danecek et al. 2021) to yield a filtered all-sites call set. Nucleotide diversity (π) was computed with scikit-allel v1.3.13 (Miles et al. 2024) as the sum of per-site mean pairwise differences over biallelic SNPs, divided by the number of callable sites (positions with genotypes called in ≥3 of 4 samples). Feature-stratified π was computed by restricting the all-sites VCF to BED intervals derived from the gene annotation (gene, exon, CDS, intron, 5′ UTR, 3′ UTR) and from the complement of the gene span against the callable genome (intergenic). Per-sample heterozygous-site counts were obtained with bcftools; heterozygous calls were also intersected with 100 kb non-overlapping windows using bedtools v2.31.1 to obtain per-window heterozygosity distributions. Runs of homozygosity were estimated with bcftools roh using a uniform default allele frequency (--AF-dflt 0.4, used in lieu of a population allele-frequency panel given the small sample size). Transition/transversion ratios were computed with VCFtools v0.1.15 and cross-validated with bcftools stats.

### Microsatellite marker discovery

Simple sequence repeats (SSRs) were identified in five *P. ficus* assemblies: the reference (VMBCA21f) and four whole-genome assemblies of MB21-2, MB21-3, MB21-P, MB21-Q. Raw reads were quality- and adapter-trimmed with Trimmomatic v0.36 (ILLUMINACLIP with TruSeq3-PE-2 adapters, 2:30:10; LEADING:7; TRAILING:7; SLIDINGWINDOW:10:20; MINLEN:36; Bolger et al. 2014). Trimmed reads were error-corrected with SPAdes BayesHammer (--only-error-correction) and then assembled *de novo* with SPAdes v3.13.0 (Bankevich et al. 2012) using --dataset corrected.yaml -k 77,97,117,127,137,145 --careful --only-assembler. SSR loci were detected with MISA v2.1 (Beier et al. 2017) using minimum repeat numbers of 6 for dinucleotides and 5 for tri-, tetra-, penta-, and hexanucleotide motifs, a maximum interruption length of 100 bp between compound SSRs, and mononucleotide repeats were excluded. Primer pairs flanking each SSR were designed with Primer3 v2.5.0 (Untergasser et al. 2012) on a 500 bp sequence window centered on each repeat (250 bp flank on each side; primer length 18-27 bp, optimum Tm 60°C with design range 57-63°C, GC 40-60%, product size 100-300 bp). Polymorphic markers were identified by anchoring loci across assemblies through perfect identity of both forward and reverse primer sequences: any primer pair recovered with identical sequences in all five assemblies was considered a candidate cross-validated marker and classified as polymorphic when the *in silico* repeat count differed between at least two individuals. Candidates were further filtered for primer pair quality (Primer3 pair penalty ≤ 5, melting temperature 58-62°C, F-R Tm difference ≤ 2°C, GC content 40-60%, product size 100-300 bp) and restricted to loci showing product-size variation strictly proportional to repeat count differences.

### RNA-Seq and sex-biased gene expression

Total RNA was isolated from three biological replicate pools of female insects (three insects per pool) and three biological replicate pools of male insects (four insects per pool) using the Quick-RNA Tissue/Insect Microprep Kit (Zymo Research, Irvine, CA, USA). RNA purity, quantity and quality were assessed using a NanoDrop 2000 spectrophotometer (Thermo Fisher Scientific, Hanover Park, IL, USA), a Qubit 2.0 Fluorometer with the RNA Broad Range Assay Kit (Life Technologies, Carlsbad, CA, USA), and electrophoresis respectively. RNAseq libraries were prepared using the Illumina TruSeq RNA Sample Preparation Kit v2 (Illumina, CA, USA) following Illumina’s low-throughput protocol. Quantity and quality of the final libraries were evaluated using a High Sensitivity DNA chip on a Bioanalyzer 2100 instrument (Agilent Technologies, Santa Clara, CA, USA) and sequenced in 100-bp single-end mode on an Illumina HiSeq instrument at the DNA Technologies Core Facility, University of California, Davis. Adult vine mealybugs were collected as three biological replicates each of females and males. Total RNA was extracted, mRNA-enriched, and sequenced on Illumina platforms in single-read mode (R1 reads from paired-end libraries were used). Reads were quality-trimmed Trimmomatic v0.36 (Bolger et al. 2014), then aligned to the decontaminated *P. ficus* reference genome (Ps + Hc) with HISAT2 v2.1.0 (Kim et al. 2019) using --very-sensitive -k 250, allowing up to 250 alignments per read to account for the haplotype duplication in the combined Ps + Hc assembly. Genome-coordinate alignments were converted to transcript coordinates with UBU v1.2 (sam-xlate) (Mose 2025) using a BED file of mRNA coordinates. Transcript-level abundances were estimated with Salmon v1.5.1 (Patro et al. 2017) in alignment-based mode (--libType U --numBootstraps 100 --seqBias --posBias) against the *P. ficus* transcriptome containing both Ps and Hc transcripts. One hundred bootstrap replicates were generated for downstream uncertainty estimation. Because the reference includes both primary contigs (Ps) and retained haplotigs (Hc) annotated independently, a subset of loci is represented twice in the transcriptome. Salmon’s expectation-maximization algorithm correctly distributes ambiguously mapped reads across these copies. Transcript-level quantifications were imported into R with tximport v1.30 (Soneson et al. 2016) and aggregated to the gene level using a tx2gene table; gene-level counts at duplicated loci are conservative with respect to statistical power, since reads are split between paralog-like haplotype copies. Differential expression between sexes was tested with DESeq2 v1.42 (Love et al. 2014) with sex as the design variable and female as the reference level. Genes were classified as differentially expressed at |log2FC| > 1 and Benjamini-Hochberg adjusted p < 0.05. Per-scaffold enrichment for sex-biased genes was tested with Fisher’s exact tests, comparing the proportion of F-biased and M-biased genes on each scaffold (≥10 tested genes) against the genome-wide background. Resulting p-values were corrected with Benjamini-Hochberg. GO enrichment of F-biased and M-biased gene sets was tested with one-sided Fisher’s exact tests (alternative = greater) against the universe of DE-tested genes carrying InterProScan-derived GO annotations. P-values were corrected with Benjamini-Hochberg. To identify protein domains over-represented among sex-biased candidate effectors, each InterPro accession was tested with a one-sided Fisher’s exact test for enrichment in the focal class (F-biased or M-biased) relative to all other DESeq2-tested effectors. P-values were adjusted across all tested domains by the Benjamini-Hochberg procedure.

### Phylogenomic analysis and divergence-time estimation

To place *P. ficus* in a broader phylogenetic context, we compiled a 17-taxon dataset comprising the *P. ficus* VMBCA21f genome assembled in this study, eight additional Pseudococcidae (*Planococcus citri*, GCF_950023065.1; *Pseudococcus jackbeardsleyi*, GCA_038380155.1; *Pseudococcus viburni*, GCA_033439095.1; *Pseudococcus longispinus*, GCA_900064475.1; *Maconellicoccus hirsutus*, GCA_003261595.1; *Phenacoccus solenopsis*, GCA_009761765.1; *Nipaecoccus viridis*, GCA_052327235.1; *Paracoccus marginatus*, GCA_048836485.1), three Coccidae (*Ceroplastes pseudoceriferus*, GCA_050872025.1; *Toumeyella liriodendri*, GCA_041937245.1; *Ericerus pela*, GCA_011428145.1), one Monophlebidae (*Icerya purchasi*, GCA_039619475.1), one Aphididae (*Acyrthosiphon pisum*, GCF_005508785.2), one Aleyrodidae (*Bemisia tabaci*, GCF_918797505.1), one Liviidae (*Diaphorina citri*, GCA_024506315.3), and *Drosophila melanogaster* (GCF_000001215.4) as a distant outgroup. Single-copy orthologs were identified in each genome with BUSCO v6.0.0 (Manni et al. 2021) against the hemiptera_odb12 database. Per-ortholog protein alignments were generated with MAFFT v7 (--auto; Katoh and Standley 2013), trimmed with trimAl v1.5 (-automated1; Capella-Gutiérrez et al. 2009), and concatenated with AMAS (Borowiec 2016) into a 1.6 Mb amino-acid supermatrix. Maximum-likelihood inference was performed with IQ-TREE v3 (Minh et al. 2020) under the LG+R4 substitution model, with 1,000 ultrafast bootstrap and 1,000 SH-aLRT replicates. All internal nodes received 100% SH-aLRT support and 98-100% UFBoot support. Divergence times were estimated with MCMCTree (PAML 4.9j; Yang 2007) using the approximate likelihood method (dos Reis and Yang 2011). The supermatrix was randomly subsampled to 200,000 amino-acid sites (seed 42) to reduce computational requirements. Branch lengths, gradient and Hessian were estimated under the LG amino-acid rate matrix with discrete Γ rate variation (α = 0.5, four categories). MCMCTree was then run under the independent-rates relaxed clock model (clock = 2), with a birth-death-sampling prior on internal node times (parameters 1, 1, 0.1), a Γ(2, 20) prior on the mean substitution rate, and a Γ(1, 10) prior on the rate variance. Three fossil-based node calibrations were applied: (i) the root, representing the Hemiptera-Diptera divergence, was constrained with soft bounds at 300-365 Ma based on Carboniferous insect fossils (Misof et al. 2014); (ii) the Coccoidea crown (*Icerya purchasi* + Neococcoidea) was given a minimum age of 130 Ma based on *Hodgsonicoccus patefactus* in Lebanese amber (Vea and Grimaldi 2015); (iii) the Neococcoidea crown (Coccidae + Pseudococcidae) was given a minimum age of 99 Ma based on *Kozarius perpetuus* and *Rosahendersonia prisca* in mid-Cretaceous Burmese amber (Vea and Grimaldi 2015; amber U–Pb zircon age 98.79 ± 0.62 Ma, Shi et al. 2012). Minimum-age constraints were implemented as soft lower bounds with the default 2.5% tail probability. Two independent MCMC chains were run from different random seeds with 500,000 burn-in generations followed by 25,000 samples taken every 200 generations. Cross-chain agreement was assessed by comparison of posterior means across all internal nodes; the maximum difference between chains was 1.56%. The two chains were pooled (after a further 10% within-chain burn-in) for the final posterior summary.

## Results

### Genome assembly of *Planococcus ficus* VMBCA21f

A reference genome was assembled for a single adult female *P. ficus* (sample VMBCA21f), sequenced with PacBio CLR. Four additional individuals from separate field-collected populations (MB21-2, MB21-3, MB21-P, MB21-Q) were sequenced with Illumina paired-end reads for resequencing. Six RNA-Seq libraries (three females, three males) were generated for gene annotation and sex-biased expression analysis. Haploid genome size was estimated by k-mer (k = 21) frequency analysis of the four Illumina samples with Jellyfish 2.2.7 and GenomeScope 2.0. Estimates ranged from 386 to 423 Mb, with heterozygosity between 0.19% and 0.33% and repeat content between 28% and 34% (**Table S1**). The unique sequence component was stable across samples (∼278-279 Mb), indicating that inter-sample size variation reflects differences in repetitive content rather than gene-space expansion. These estimates are broadly consistent with the ∼370 Mb primary assembly span (Ps; below).

PacBio CLR reads from a single female insect (VMBCA21f) were assembled with FALCON-Unzip, producing 1,755 primary contigs (N50 = 356 kb, 368.6 Mb) and 4,072 alternative haplotigs (N50 = 39 kb, 127.0 Mb). After Arrow polishing, the primary contigs were scaffolded with SSPACE, reducing the number of sequences from 1,755 to 1,323 and increasing the N50 from 356 to 485 kb; haplotigs were retained as Arrow-polished contigs without further scaffolding. The assembly was screened for non-host sequence by per-scaffold blastn against a custom local database of reference genomes spanning bacteria, archaea, fungi, plants, and metazoans. Four sequences, three primary scaffolds and one haplotig (573 kb, 0.12% of the assembly), were identified as contamination and removed: two showed end-to-end identity to *Candidatus Moranella endobia* (NC_021057.1, NC_015735.1; query coverage 87-95%, identity >92%), the γ-proteobacterial endosymbiont nested inside *Tremblaya princeps* in Pseudococcidae, while two additional scaffolds were identified as fungal contamination.

The decontaminated v1 reference comprises a primary assembly (Ps) of 1,320 scaffolds spanning 369.4 Mb (N50 = 484.9 kb, L50 = 239), and an associated set of 4,071 alternative haplotigs (Hc) spanning 127.0 Mb (N50 = 39.1 kb; **Table 1**, **Table S2**). The Ps assembly contains 40 scaffolds longer than 1 Mb (largest 2.68 Mb) and 931 scaffolds longer than 100 kb. GC content is uniform across partitions (Ps 34.5%, Hc 34.6%); gaps comprise 0.37% of Ps and are absent from Hc. Whole-genome alignment of the *P. ficus* assembly to the chromosome-level *P. citri* reference (GCF_950023065.1; Ross et al. 2024) supported the structural integrity of the assembly: 240 Mb of *P. ficus* sequence (65% of the assembly) aligned to the five *P. citri* chromosomes via minimap2, and gene-level synteny analysis with MCScanX recovered 619 cross-species collinear blocks containing 9,123 orthologous gene pairs (Beier et al. 2017; Tang et al. 2008; Wang et al. 2012). The 532 large *P. ficus* scaffolds (≥100 kb aligned) mapped uniquely and contiguously to one of the five *P. citri* chromosomes (**Figure S1**), with no evidence of inter-chromosomal mis-joins, consistent with conserved macrosynteny across ∼2.5 Ma of *Planococcus* divergence.

**Table 1.** Reference assembly statistics for *P. ficus* VMBCA21f.

| Metric | Primary (Ps) | Haplotigs (Hc) |
| --- | --- | --- |
| Scaffolds | 1,320 | 4,071 |
| Total length (Mb) | 369.4 | 127.0 |
| Largest scaffold (kb) | 2,677.8 | 307.0 |
| Scaffold N50 (kb) | 484.9 | 39.1 |
| L50 | 239 | 912 |
| GC content (%) | 34.5 | 34.6 |
| Gap content (%) | 0.37 | 0.00 |
| Repeat-masked (%) | 32.0 | 34.4 |
| Genes | 23,489 | 7,591 |
| mRNAs | 33,259 | 10,374 |
| BUSCO genome C/F/M (%) | 90.0 / 1.7 / 8.3 | 28.9 / 3.3 / 67.8 |
| BUSCO protein C/F/M (%) | 86.1 / 2.9 / 11.0 | 25.9 / 3.3 / 70.8 |

Gene prediction was performed on the decontaminated assembly with a combined evidence pipeline integrating *ab initio* predictions, spliced alignment of RNA-Seq evidence assembled with Trinity and StringTie, and protein homology, consolidated by EVidenceModeler and refined by PASA. The Ps annotation contains 23,489 protein-coding genes producing 33,259 transcripts (mean 1.42 isoforms per gene); the Hc annotation contains 7,591 genes and 10,374 transcripts (1.37 isoforms per gene) (**Table 1**). Repeat annotation identified 549,560 features covering 118.0 Mb of non-overlapping sequence on Ps (32.0%) and 204,313 features covering 43.7 Mb on Hc (34.4%; **Table S3**). Both partitions are dominated by unclassified repeats (Ps 21.4%, Hc 22.8%), followed by DNA transposons (Ps 5.5%, Hc 5.9%), LTR retroelements (Ps 3.0%, Hc 3.3%), and rolling-circle Helitron elements (Ps 1.8%, Hc 2.1%); LINEs, SINEs, satellites, and simple/low-complexity sequence together contribute the remaining ∼2.4% in each partition (**Table S3**). The high proportion of unclassified repeats reflects the limited prior characterization of repeat families in Pseudococcidae and the use of a *de novo* repeat library generated for this assembly.

Completeness was assessed with BUSCO v6.0.0 against the hemiptera_odb12 lineage (n = 3,396; **Table S4**). The primary assembly recovered 90.0% complete BUSCOs in genome mode and 86.1% in protein mode (longest isoform per gene). The haplotigs recovered substantially fewer markers in either mode (28.9% genome, 25.9% protein), consistent with their partial coverage of the genome as alternative haplotypes.

### The *P. ficus* candidate effector repertoire

From the 23,489-protein *P. ficus* primary-scaffold proteome (longest isoform per gene), SignalP 6.0 predicted a signal peptide for 3,062 proteins; 2,901 of these had an SP probability ≥ 0.7. Restricting to a mature peptide length of 30-500 amino acids retained 2,129 candidate effectors (9.1% of the proteome) for downstream analyses (**Table S5**). The mature peptides spanned a wide size range (median 248 aa, IQR 156-364). Cysteine content was modest overall (median 1.64%, IQR 0.68-2.91%), with only 333 candidates (15.6%) reaching the ≥ 4% cysteine threshold conventionally used to delimit the classical disulfide-stabilized small Cys-rich effector class. The *P. ficus* candidate effector repertoire is therefore dominated by proteins outside this canonical category.

InterProScan recovered at least one InterPro domain in 1,264 of the 2,129 candidates (**Fig. 2A**; 59.4%); the remaining 865 (40.6%) had no detectable InterPro signature, indicating a large reservoir of small, possibly fast-evolving secreted proteins whose function cannot be inferred from sequence alone. Within the annotated subset, the most abundant functional groups (**Fig. 2B**; **Table 2**) fell into three broad groups. Hydrolytic and digestive enzymes were prominent, including S1 chymotrypsin/trypsin-like serine proteases (n = 50), C1A papain-like cysteine peptidases of the cathepsin family (n = 43), α/β hydrolase-fold enzymes (n = 64), triacylglycerol lipases (n = 27), UDP-glycosyltransferases (n = 45), and carbonic anhydrases (n = 32); a serpin family (n = 30) likely provides negative regulation of the serine protease component. Binding and recognition proteins formed a second group, headed by immunoglobulin-like-fold proteins (n = 76), pheromone/general odorant-binding proteins (n = 60), PEBP-like proteins (n = 46), annexins (n = 26), and juvenile-hormone-binding/Takeout proteins (n = 21). A third group of cuticle-associated proteins (n = 43) and leucine-rich repeat proteins (n = 34) accounted for additional structural and recognition functions. The composition is consistent with the biology of *P. ficus* as a phloem-feeding insect that secretes a complex hydrolytic and regulatory arsenal during feeding, alongside chemoreceptive and developmental components.

**Figure 2.**
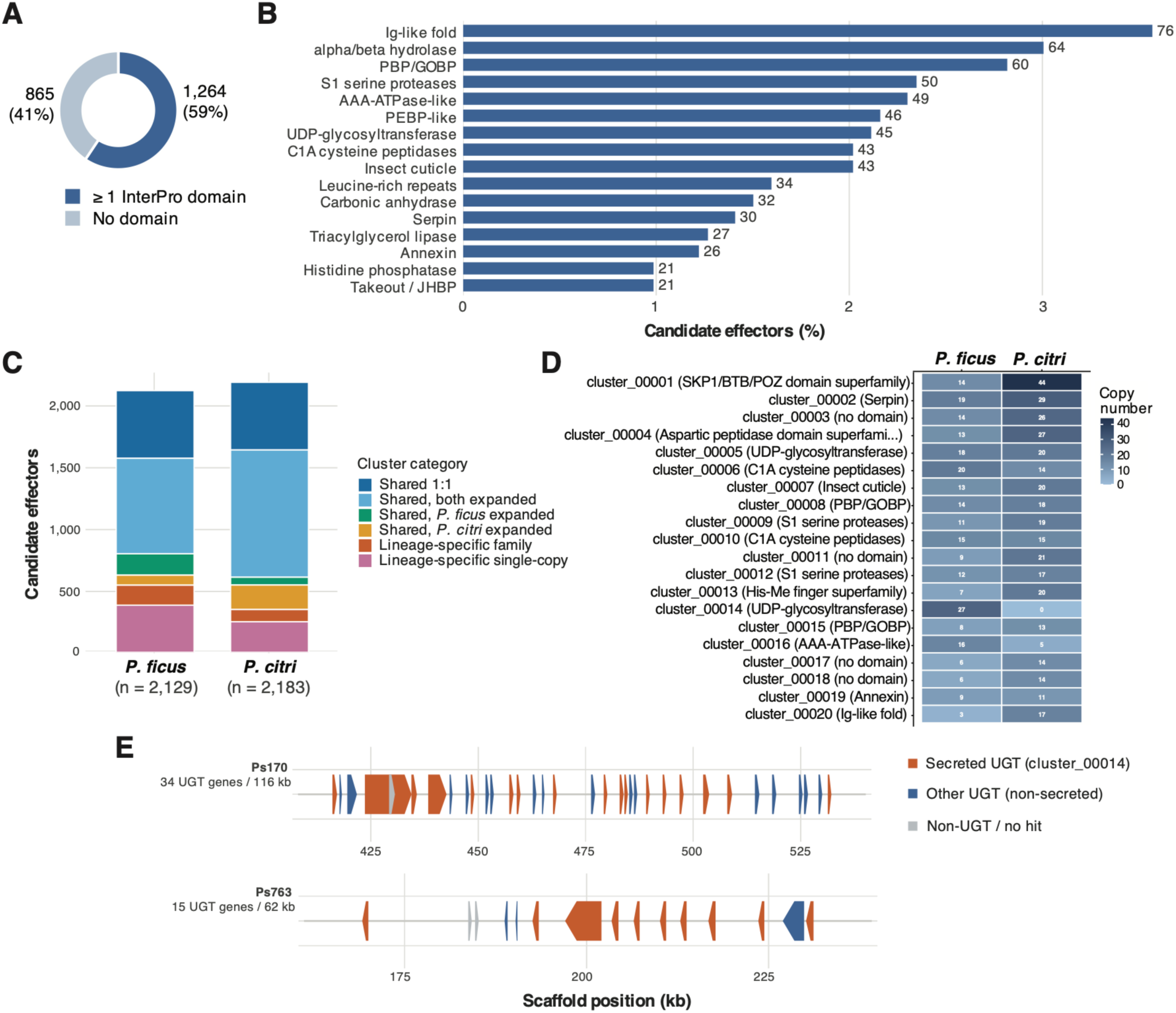
Candidate effector repertoire of *Planococcus ficus* and cross-species comparison with *P. citri.* **(A)** InterPro domain annotation coverage of the 2,129 candidates. **(B)** Major InterPro functional groups among the 2,129 *P. ficus* candidate effectors, shown as the percentage of candidates carrying at least one domain in each group (counts above bars; groups defined by collapsing related InterPro accessions, see **Table 2**). **(C)** Composition of the jointly clustered secretomes of *P. ficus* (n = 2,129) and *P. citri* (n = 2,183), partitioned by cluster category: shared single-copy 1:1 orthologues, shared clusters expanded in both or one lineage, and lineage-specific clusters (multi-copy families or single-copy). **(D)** The 20 most populous effector clusters, showing per-species copy number (cell values; colour scale at right). Dominant InterPro annotation per cluster in parentheses. Copy numbers are of secreted (candidate-effector) members only. **(E)** Genomic organization of the *P. ficus*-specific UDP-glycosyltransferase expansion (cluster_00014). Two tandem arrays, on scaffolds Ps170 (34 UGT genes, ∼116 kb) and Ps763 (15 UGT genes, ∼62 kb), are drawn to scale with genes as true-width rectangles and a terminal tick indicating strand. The 27 secreted UGTs that constitute cluster_00014 (orange) are interspersed with non-secreted UGTs of the same family (blue); three short non-UGT or unannotated gene models are shown in grey. The arrays are absent from *P. citri*. The 27 in panel D therefore refers to the secreted subset of these 49-gene arrays.

**Table 2.** Major InterPro functional groups in *P. ficus* candidate effectors. Functional groups defined by collapsing related InterPro accessions within the same family or superfamily. Counts are unique proteins per group among the 2,129 *P. ficus* candidate effectors. Groups are ordered by decreasing abundance.

| Functional group | Likely role | Unique proteins | % of 2,129 |
| --- | --- | --- | --- |
| Immunoglobulin-like fold (Ig-like superfamily) | Cell adhesion, recognition, binding | 76 | 3.57 |
| $\alpha/\beta$ hydrolase fold | Broad hydrolase scaffold (esterases, lipases, peptidases) | 64 | 3.01 |
| Pheromone/general odorant-binding protein (PBP/GOBP) | Chemoreception, small-molecule transport | 60 | 2.82 |
| S1 serine proteases (chymotrypsin/trypsin-like) | Extracellular proteolysis, digestion | 50 | 2.35 |
| AAA-ATPase-like | ATP-dependent remodeling/chaperone activity | 49 | 2.30 |
| PEBP-like | Phosphatidylethanolamine binding; signaling regulation | 46 | 2.16 |
| UDP-glycosyltransferases | Glycosylation of small molecules; xenobiotic detoxification | 45 | 2.11 |
| C1A papain-like cysteine peptidases (cathepsins) | Lysosomal/extracellular protein degradation | 43 | 2.02 |
| Insect cuticle proteins | Cuticle structure and assembly (R&R-type chitin-binding) | 43 | 2.02 |
| Leucine-rich repeats | Protein-protein interaction, immune recognition | 34 | 1.60 |
| Carbonic anhydrases | CO<sub>2</sub>/HCO<sub>3</sub><sup>-</sup> interconversion; pH and ion homeostasis | 32 | 1.50 |
| Serpins (serine protease inhibitors) | Regulation of protease cascades; immunity | 30 | 1.41 |
| Triacylglycerol lipases | Lipid hydrolysis | 27 | 1.27 |
| Annexins | Calcium/phospholipid binding; membrane trafficking | 26 | 1.22 |
| Histidine phosphatases | Phosphate metabolism | 21 | 0.99 |
| Juvenile-hormone binding (Takeout/JHBP) | Hormone transport; insect development and behavior | 21 | 0.99 |

The combined *P. ficus* and *P. citri* secretomes were jointly clustered with MCL to assess effector conservation between the two species. The 4,312 candidate effectors partitioned into 1,596 clusters (**Fig. 2C**, **Tables S7** and **S8**). Most effectors of both species belonged to shared clusters with cross-species homologues: 74% of *P. ficus* effectors (1,573 / 2,129) and 84% of *P. citri* effectors (1,837 / 2,183; **Table S6**) were assigned to a cluster containing members of both species. Shared single-copy orthologues (547 clusters; 1:1 between the two species) represented the deepest core of conserved effectors. Lineage-specific clusters were also abundant, but their counts were asymmetric: *P. ficus* had 556 lineage-specific effectors (381 singletons + 175 in family expansions), compared with only 346 in *P. citri* (247 + 99). The two species therefore share a substantial conserved effector core but differ in the size of their lineage-specific repertoires, with *P. ficus* harbouring more novel and expanded private effector families.

The 20 most populous clusters together accounted for 618 effector proteins (14.3% of the candidate set) and were dominated by a small number of recurring functional categories (**Fig. 2D**, **Table S7**). Detoxification and digestive enzymes were prominent: two clusters of UDP-glycosyltransferases (clusters 5 and 14), two clusters of C1A papain-like cysteine peptidases (clusters 6 and 10), and two clusters of S1 serine proteases (clusters 9 and 12) together comprised six of the top 20 expansions. Protease regulation appeared as the second most populous cluster overall (cluster_00002, a 48-member serpin family). Chemoreception was represented by two pheromone/general odorant-binding protein expansions (clusters 8 and 15). The largest single cluster (cluster_00001, 58 members) was annotated as SKP1/BTB/POZ domain proteins, which most commonly serve as ubiquitin-ligase adaptors; whether the cluster represents a bona fide secreted variant or a misclassified intracellular family in both genomes requires further validation.

Of the 20 most populous clusters, four had no detectable InterPro domain (clusters 3, 11, 17, 18; jointly 110 proteins), representing a substantial reservoir of conserved but functionally uncharacterized cross-species effector families. Several clusters showed pronounced asymmetric expansion between species. cluster_00014 (UDP-glycosyltransferase) is entirely *P. ficus*-specific, with 27 secreted members and no detectable homologue in *P. citri*. These 27 secreted UGTs are the signal-peptide-bearing subset of two larger *P. ficus*-specific UDP-glycosyltransferase tandem arrays, comprising 34 genes on scaffold Ps170 (∼116 kb) and 15 on Ps763 (∼62 kb); within each array the secreted and non-secreted UGTs are interspersed and uniformly oriented, consistent with tandem duplication rather than an assembly artefact (**Fig. 2E**). cluster_00016 (AAA-ATPase-like; 16 Pfic vs 5 Pcit) represents a more modest *P. ficus*-biased expansion. Conversely, cluster_00013 (His-Me finger nuclease superfamily; 7 Pfic vs 20 Pcit) and cluster_00020 (Ig-like fold; 3 Pfic vs 17 Pcit) are expanded in *P. citri*. These lineage-specific expansions likely reflect divergent host-adaptation pressures between the two species, with the *P. ficus*-specific UGT expansion of particular interest given the role of UGTs in xenobiotic detoxification of plant secondary metabolites.

### Diversity and microsatellite discovery

The four insects (MB21-2, MB21-3, MB21-P, MB21-Q), collected from four vineyards in California’s San Joaquin Valley, were sequenced to depths of 65 to 82 million read pairs each (151 bp paired-end), yielding 49 to 61x mean coverage of the primary assembly. Mapping rates were consistently high across all four samples, ranging from 95.8% to 96.3% mapped reads, with 87.8% to 90.3% in proper pairs (**Table S9**). Joint genotyping produced 2,247,614 raw variants. After excluding three contaminating scaffolds and applying GATK hard filters, the final call set comprised 1,232,504 SNPs and 517,359 indels (PASS rates 73.6% and 92.6%, respectively). Per-individual coverage at variant sites ranged from 35.8x to 46.7x. Each individual carried 558,223-682,850 heterozygous sites, with a mean of 127-156 heterozygous sites per 100 kb window. Genome-wide nucleotide diversity, estimated from the all-sites VCF using callable-site denominators, was π = 2.06 x 10⁻³ over 337.2 Mb of callable genome.

Feature-stratified analysis showed that intergenic regions had the highest π (2.09 x 10⁻³), followed by introns (2.03 x 10⁻³) and coding sequence (2.02 x 10⁻³), with the two UTR classes substantially depleted (5′ UTR: 1.40 x 10⁻³; 3′ UTR: 1.41 x 10⁻³; **Table S10**). SNP density per kb showed a similar UTR depletion (5′ UTR: 2.33; 3′ UTR: 2.11) but was elevated in CDS (3.75) relative to intergenic (3.33); this elevation is attributable to reduced mappability in repeat-rich intergenic sequence and is not seen in π estimated against the callable denominator. The Ts/Tv ratio was 1.72 genome-wide, rose to 2.37 in coding sequence, and was lower in introns (1.63) and intergenic regions (1.69), consistent with selection against non-synonymous transversions in coding regions. Ts/Tv was stable across allele-count classes (1.68-1.78), across variant-quality bins, and outside repeat-annotated regions (1.76). bcftools-derived Ts/Tv (1.70-1.71) agreed with VCFtools to within 0.02 (**Table S11**).

To enable marker development for population studies, each of the four individuals was additionally assembled *de novo* with SPAdes and screened for simple sequence repeats alongside the chromosome-level reference. MISA recovered 21,088-21,384 perfect SSR loci per assembly (**Table S12**). Cross-assembly anchoring by exact forward and reverse primer identity yielded 953 loci independently recovered in all five assemblies: 718 were monomorphic and 235 showed repeat-count variation between individuals. Filtering for primer quality and strict repeat-proportional product-size variation retained 132 cross-validated polymorphic SSR markers (**Table S13**).

### Pervasive sex-biased expression

Sex-biased expression was pervasive in *P. ficus*. Among the 18,013 Ps host genes tested with DESeq2, 7,362 (40.9%) were significantly differentially expressed between sexes (|log₂FC| > 1, BH-adjusted P < 0.05; **Table S14**): 4,500 F-biased and 2,862 M-biased on the Ps partition (**Fig. 3A, B**), with a comparable but smaller bias on the haplotig partition (1,578 F-biased, 876 M-biased; **Fig. S2 A, B**). At the genome-wide level the median log₂FC across all expressed Ps genes was −0.33, shifted toward female bias but falling short of the −1 expected under naive halving of paternal alleles in males (**Fig. 3C**). This is consistent with the incomplete somatic silencing reported in *P. citri* (de la Filia et al. 2021), though bulk RNA-seq cannot distinguish residual paternal expression from active dosage compensation.

**Figure 3.**
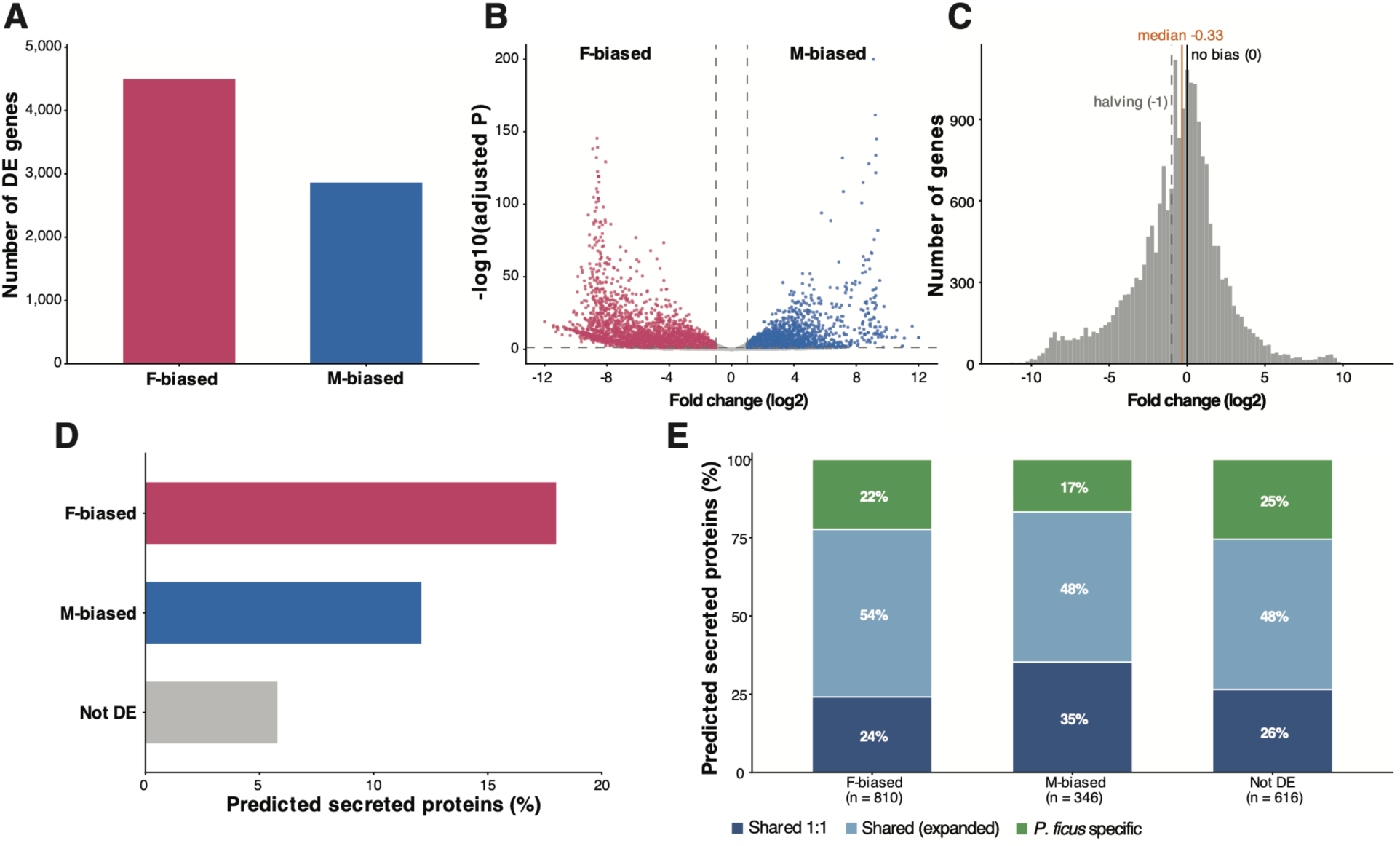
Sex-biased gene expression and the secreted-protein repertoire in *P. ficus*. Differential expression between adult males and females across the primary-scaffold (Ps) gene set, with female as the reference (log₂ fold change = M/F); sex-biased at |log₂FC| > 1 and adjusted P < 0.05 (Benjamini-Hochberg). **(A)** Counts of sex-biased Ps genes by direction. **(B)** Volcano plot of log₂FC against significance; female-biased (pink), male-biased (blue), not differentially expressed (grey); dashed lines mark the |log₂FC| = 1 and P = 0.05 thresholds. **(C)** Distribution of log₂FC across all Ps loci, with the observed median (orange), no-bias expectation (0, black), and naive paternal-halving prediction (−1, dashed grey) marked. **(D)** Predicted secreted proteins (SignalP 6.0, SP probability ≥ 0.7, mature peptide 30-500 aa) as a percentage of Ps genes in each DE class. **(E)** Candidate effectors by DE class and cross-species conservation, scored against *Pseudococcus citri* orthologous clusters as Shared 1:1, Shared with family expansion, or *P. ficus*-specific; n below each bar is the number of candidate effectors with cluster assignments.

Functional enrichment (**Fig. S3, Table S15**) identified distinct sex-specific programs. Male-biased genes were enriched for the locomotor and sensory machinery of dispersing adult males: odorant binding (n = 49, fold enrichment 3.1, padj = 1.1 x 10⁻¹³, the strongest male-biased term), nervous system process (n = 120, 2.0x), muscle system process, cilium and synapse organization, and ion transport. Female-biased genes were enriched for translation and ribosome biogenesis (n = 203, 2.4x, padj = 1.4 x 10⁻³⁸, the most significant term overall), mRNA metabolism, and nuclear/nucleolar components, alongside more specific categories including negative regulation of the ERK1/2 cascade (4.5x), carboxylic ester hydrolase activity (n = 107, 2.7x), antimicrobial defense, DNA repair, and gene silencing by RNA.

Given the female-specific feeding behavior of *P. ficus*, we asked whether the candidate secreted proteins identified above showed a corresponding female bias. F-biased genes were strongly enriched for predicted secretion (18.0%) compared with M-biased genes (12.1%) and the non-differentially-expressed background (5.8%); the F-biased Fisher’s odds ratio of 2.86 (P = 3 x 10⁻⁸⁹) was over twice that of M-biased (OR 1.32, P = 1.6 x 10⁻⁵; **Fig. 3D**). Stratifying these candidate effectors by their cross-species cluster category revealed that F-biased effectors tracked the global *P. ficus* effector distribution, whereas M-biased effectors were significantly enriched in deeply conserved shared 1:1 orthologs with *P. citri* relative to F-biased effectors (35% vs 24%; Fisher’s exact test, OR 1.72, P = 1.3 x 10⁻⁴; **Fig. 3E**). The two sex-biased secretomes therefore differ in the conservation of their members: the male-biased set is drawn preferentially from an ancestral, conserved repertoire, while the female-biased set spans the more lineage-specific *P. ficus* effector pool.

Among M-biased effectors, eight InterPro domains were significantly enriched (**Table 3**), with two clear functional themes. Male-specific chemoreception was the dominant signal: pheromone/general odorant-binding proteins (PBP/GOBP; IPR036728 and IPR006170) reached the strongest enrichment of the analysis (OR 6.2 and 5.7, BH-adjusted P = 9.2 x 10⁻⁸ and 6.6 x 10⁻⁶). Adult cuticle and exoskeleton biology was the second: three insect cuticle protein families (IPR000618, IPR050468, IPR051217) and the chitin-binding R&R consensus motif (IPR031311) were all enriched among M-biased effectors. Two additional families with putative neural/developmental roles (sleep homeostasis regulator IPR050975; UPAR/Ly6 IPR031424) rounded out the M-biased signature. These functions match the biology of adult *P. ficus* males, which are short-lived, winged, non-feeding morphs whose primary roles are mate location and reproduction.

**Table 3.** InterPro domains significantly enriched in sex-biased *P. ficus* candidate effectors. One-sided Fisher’s exact test for enrichment in the focal DE class relative to the remaining tested *P. ficus* effectors; *P*-values adjusted by Benjamini-Hochberg. Only domains with BH-adjusted *P* < 0.05 are shown (OR = odds ratio).

| DE class | InterPro accession | Description | % focal | % background | OR | BH-adj $P$ |
| --- | --- | --- | --- | --- | --- | --- |
| M-biased | IPR036728 | Pheromone/general odorant-binding protein domain superfamily | 9.25 | 1.61 | 6.21 | $9.2 \times 10^{-8}$ |
| | IPR006170 | Pheromone/general odorant-binding protein | 7.51 | 1.40 | 5.70 | $6.6 \times 10^{-6}$ |
| | IPR000618 | Insect cuticle protein | 6.36 | 1.19 | 5.62 | $6.1 \times 10^{-5}$ |
| | IPR031311 | Chitin-binding type R&R consensus | 4.05 | 0.63 | 6.63 | $2.4 \times 10^{-3}$ |
| | IPR050975 | Sleep homeostasis regulator | 2.31 | 0.21 | 11.16 | $2.6 \times 10^{-2}$ |
| | IPR031424 | UPAR/Ly6 domain-containing protein | 2.60 | 0.35 | 7.58 | $3.4 \times 10^{-2}$ |
| | IPR051217 | Insect cuticle structural | 2.89 | 0.49 | 6.02 | $3.6 \times 10^{-2}$ |
| | IPR050468 | Larval/pupal cuticle protein | 3.18 | 0.63 | 5.16 | $3.6 \times 10^{-2}$ |

| DE class | InterPro accession | Description | % focal | % background | OR | BH-adj *P* |
| --- | --- | --- | --- | --- | --- | --- |
| F-biased | IPR036610 | PEBP-like superfamily | 5.19 | 0.42 | 13.06 | 3.4 x 10<sup>-8</sup> |
|  | IPR035810 | Phosphatidylethanolamine-binding protein | 3.46 | 0.31 | 11.40 | 6.7 x 10<sup>-5</sup> |
|  | IPR018631 | AAA-ATPase-like domain | 4.69 | 0.94 | 5.21 | 1.4 x 10<sup>-4</sup> |

In contrast, only three InterPro domains were significantly enriched among F-biased effectors (**Table 3**), members of just two families: the phosphatidylethanolamine-binding/PEBP-like family (IPR036610, IPR035810; OR 13.1 and 11.4) and an AAA-ATPase-like domain (IPR018631). The much smaller number of enriched domains in the F-biased set, despite F-biased effectors being more numerous than M-biased (810 vs 346 candidates), indicates that the female feeding secretome is functionally heterogeneous and not dominated by a single conserved domain architecture, consistent with the absence of a classical cysteine-rich signature reported above. A substantial fraction of F-biased effectors carries no detectable InterPro domain and likely represents novel, *P. ficus*-specific secreted proteins awaiting structural and functional characterization.

### Genomic clustering of sex-biased genes

To identify genomic regions enriched for sex-biased expression, Ps scaffolds with at least 10 tested genes (n = 260) were screened by Fisher’s exact test (BH-corrected). Six scaffolds showed significant enrichment (padj < 0.05; **Figure 4**, **Table 4**), ranging from 145 kb to 824 kb in size: five enriched for female-biased genes (Ps350, 439 kb; Ps72, 824 kb; Ps138, 549 kb; Ps376, 569 kb; Ps743, 145 kb) and one for male-biased genes (Ps328, 378 kb). The most extreme cases were Ps350 (66 genes on the scaffold, 58 tested, of which 46 sex-biased: 45 F-biased, 1 M-biased; padj = 3.2 x 10⁻⁶) and Ps72 (103 genes, 81 tested, of which 37 sex-biased: 36 F-biased, 1 M-biased; padj = 1.4 x 10⁻⁴). To test whether this clustering exceeded the level expected by chance, we permuted sex-bias labels across all tested Ps genes 1,000 times and recomputed scaffold enrichment with the same procedure. The permutation null produced a mean of 0.03 significantly enriched scaffolds (range 0-2) for each direction, far below the observed counts (empirical P < 0.001). Sex-biased expression in *P. ficus* is therefore not only pervasive but also strongly segregated into discrete genomic blocks. Directional tests (separate one-sided Fisher tests for female and male enrichment per scaffold) identified six additional scaffolds (130-554 kb) with extreme one-sided enrichment (three F-only: Ps233, Ps353, Ps799; three M-only: Ps130, Ps236, Ps259; **Table S16**), all consistent with the segregation pattern (**Fig. S4**).

**Figure 4.**
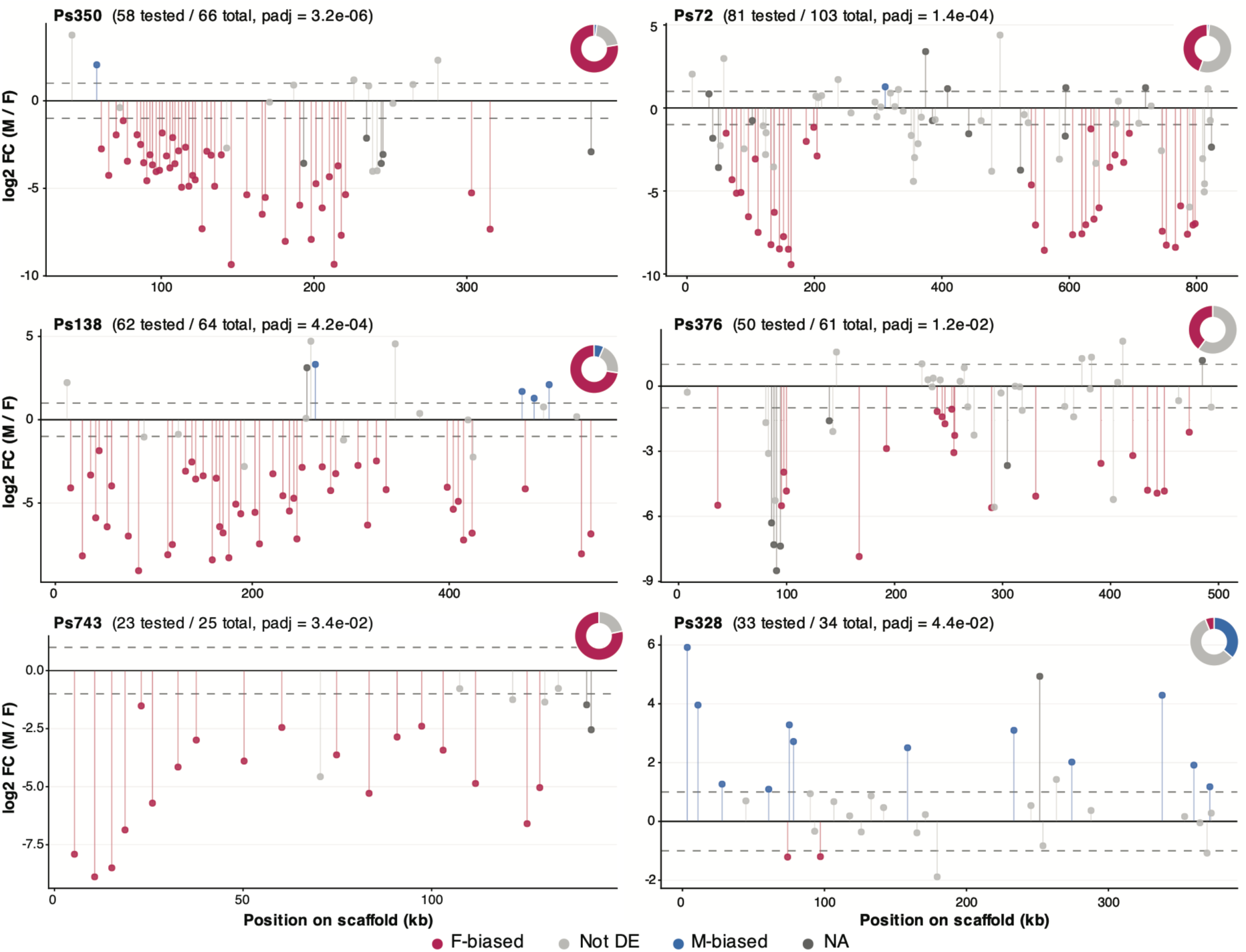
Significant sex-biased *P. ficus* scaffolds. Differential expression along the six Ps scaffolds that are significantly enriched for sex-biased genes (directional Fisher’s exact test, BH-adjusted P < 0.05). Each panel shows per-gene log₂ fold change (male / female) plotted against the gene’s midpoint position on the scaffold; colors indicate DE class (F-biased, log₂FC < −1 and padj < 0.05; M-biased, log₂FC > 1 and padj < 0.05; Not DE; or NA for genes not tested by DESeq2). Vertical segments connect each point to the zero baseline. Panel titles report the scaffold-level Fisher-test adjusted P-value. The donut inset in each panel summarizes the F-biased / Not DE / M-biased composition among tested genes on that scaffold.

**Table 4.** Primary-assembly scaffolds enriched for sex-biased gene expression in *P. ficus*. Six Ps scaffolds (≥10 DE-tested genes) showed significant enrichment for female- or male-biased genes by Fisher’s exact test (BH-corrected padj < 0.05). Size, scaffold length in kilobases; Genes, number of DE-tested loci on the scaffold; M and F, number of male- and female-biased genes (|log₂FC| > 1, padj < 0.05); Direction, dominant sex-bias category.

| Scaffold | Size (kb) | Genes | M | F | padj | Direction |
| --- | --- | --- | --- | --- | --- | --- |
| Pficus_VMBCA21f_v1.0_Ps350 | 439 | 58 | 1 | 45 | $3.2 \times 10^{-6}$ | F-biased |
| Pficus_VMBCA21f_v1.0_Ps72 | 824 | 81 | 1 | 36 | $1.4 \times 10^{-4}$ | F-biased |
| Pficus_VMBCA21f_v1.0_Ps138 | 549 | 62 | 4 | 45 | $4.2 \times 10^{-4}$ | F-biased |
| Pficus_VMBCA21f_v1.0_Ps376 | 569 | 50 | 0 | 20 | $1.2 \times 10^{-2}$ | F-biased |
| Pficus_VMBCA21f_v1.0_Ps743 | 145 | 23 | 0 | 18 | $3.4 \times 10^{-2}$ | F-biased |
| Pficus_VMBCA21f_v1.0_Ps328 | 378 | 33 | 12 | 2 | $4.4 \times 10^{-2}$ | M-biased |

Inspection of gene content within the F-enriched scaffolds revealed clustered gene-family expansions, with each F-block representing a dense local concentration of a paralogous family that is also dispersed across other scaffolds. Ps350 contains a cluster of >10 phosphatidylethanolamine-binding proteins (PEBPs, including PEBP-like, protein D2, and protein D3 paralogs), accounting for 9 of the 18 F-biased genes contributing to the “negative regulation of ERK1/2 cascade” GO term reported above (50%), with most of the remainder on the syntenic haplotigs Hc000358F_001 and Hc000358F_003. Ps72 carries a cluster of glycosyl- and sulfo-transferases (galactose-3-O-sulfotransferases, β1,4-galactosyltransferase 7, galactosylceramide sulfotransferases) representing the largest concentration of this family in the genome (11 of 43 F-biased members), with an additional 6 paralogs on the F-only supplementary scaffold Ps799 and 7 distributed across syntenic Hc000072F haplotig fragments. Ps743 (145 kb, 14 F-biased carboxylesterase paralogs) is the densest cluster of F-biased carboxylesterases in the genome, although additional F-biased members of this family are dispersed across multiple Ps scaffolds. The single main-text M-enriched scaffold, Ps328 (34 genes on the scaffold, 33 tested, of which 14 sex-biased: 12 M-biased, 2 F-biased; padj = 4.4 x 10⁻²), was functionally heterogeneous and included a copia-like retrotransposon, a cytochrome P450 (CYP4g15-like), arylalkylamine N-acetyltransferase, fibroblast growth factor receptor-like, and synaptotagmin-14; the three additional M-only scaffolds (Ps130, Ps236, Ps259) similarly carried zinc finger paralogs, retrovirus-related Pol polyproteins, and various signaling and transmembrane components rather than a single coherent gene family.

To characterize these blocks further, we examined three additional properties of the enriched scaffolds: gene density, effect size, and within-block co-expression (**Table S17**). F-enriched scaffolds were significantly more gene-dense than the genome-wide median (0.121 vs 0.059 genes/kb; Wilcoxon P = 2.5 x 10⁻⁴), consistent with the clustered gene-family expansions described above; M-enriched scaffolds were not significantly denser than other scaffolds (P = 0.18). Effect sizes were larger for F-biased genes residing within F-blocks than for F-biased genes elsewhere (median |log₂FC| 4.93 vs 3.91; Wilcoxon P = 1.0 x 10⁻⁶); the same comparison was not significant for M-biased genes (P = 0.74). Within each block, the genes contributing to the sex bias were highly co-expressed across samples: the median pairwise Spearman correlation among sex-biased genes within each enriched scaffold ranged from 0.71 to 0.89, far above the null distribution generated by 500 random 20-gene sets drawn from genome-wide sex-biased genes (mean ρ = 0.10, 95th percentile 0.71; 11 of 12 enriched scaffolds significant at null P ≤ 0.004; **Table S18**). Repeat content of enriched scaffolds did not differ significantly from the genome-wide distribution (F-blocks median 33.9% repetitive sequence, M-blocks 31.3%, other scaffolds 37.7%; Wilcoxon P ≥ 0.15 for both comparisons), and neither retrotransposon nor DNA-element fractions were elevated in either block class. Together, these patterns indicate that sex-biased blocks correspond to coordinately co-expressed genomic regions rather than to repetitive sequence compartments, with the F-blocks additionally enriched for clustered tandem paralogs.

## Discussion

### A reference genome for *Planococcus ficus*

VMBCA21f is the first long-read, scaffolded, partially diploid reference genome for *P. ficus*, following an earlier short-read draft that is not publicly available (Corcoran and Mahaffee 2024). The 369.4 Mb primary assembly is 90.0% complete in genome-mode BUSCO and carries 23,489 annotated genes, with size, GC content, and repeat fraction matching k-mer estimates from four California samples and within the Pseudococcidae range (M. Li et al. 2020; Ross et al. 2024). FALCON-Unzip retained a further 127.0 Mb of haplotigs representing alternate alleles at heterozygous regions. This assembly has two limitations: it is not chromosome-scale, and a minority of complete BUSCOs (271 of 3,058) carry internal stop codons, likely reflecting residual PacBio CLR indel error.

Alignment to *P. citri* corroborates the large-scale structure of the assembly (**Fig. S1**). This high collinearity reflects the close relationship between the two species, whose divergence we date to 2.49 Ma (95% HPD 1.71–3.37 Ma; **Fig. 1**), the first molecular estimate for this split. At deeper nodes, the Pseudococcidae crown, the Phenacoccinae / Pseudococcinae split (Hardy et al. 2008; Choi and Lee 2022), dates to 69 Ma (95% HPD 55–84 Ma). This is younger than the genome-scale estimate of Deng et al. (2025), who dated the family to 128.1–93.2 Ma, and younger still relative to the 210–165 Ma neococcoid origin of Vea and Grimaldi (2016), consistent with Pseudococcidae being a comparatively recent radiation within neococcoids. This difference is largely one of calibration. Deng et al. placed a mid-Cretaceous fossil minimum, *Williamsicoccus megalops*, directly on the family. We excluded this fossil: it is known only from a male specimen whose placement within Pseudococcidae is unresolved, and Vea and Grimaldi (2016) likewise treated it as a floating tip rather than a node minimum. Our nearest constraint is therefore the deeper Neococcoidea node, so the 69 Ma estimate carries no direct fossil minimum and may be a lower bound.

### Candidate effector repertoire

The *P. ficus* secretome encompasses proteins released into multiple extracellular compartments, including the gut lumen during digestion and detoxification of ingested plant material, the hemolymph, and saliva injected into host tissues during stylet penetration. The salivary fraction is of particular interest for plant-insect interactions: in other phloem-feeding hemipterans these secretions suppress sieve element occlusion, modulate plant defense signaling, and in some cases mediate virus transmission (Tsai et al. 2008; Will et al. 2013). SignalP-based screens detect the signal peptide common to all compartments and cannot separate salivary from non-salivary secreted proteins without tissue-specific expression data, which is not yet available for *P. ficus*. We screened the *P. ficus* primary proteome with SignalP 6.0 as a starting point for the full secretome and compared it with that of *P. citri*, the closest sequenced relative. Although both species are polyphagous, *P. citri* has a substantially broader host range than *P. ficus* (>70 *vs*. >16 recorded plant families; García-Morales et al. 2016), and the comparison can expose lineage-specific changes in the feeding-related secretome. The two secretomes are similar in size (2,129 vs 2,183 candidates) and share a substantial conserved core, with roughly three-quarters of *P. ficus* and a slightly higher fraction of *P. citri* candidates falling into cross-species clusters. This shared core is dominated by small proteins with no detectable InterPro domain, together with conserved binding and recognition families (PEBP-like, odorant-binding, immunoglobulin-like, and juvenile-hormone-binding proteins), while enzyme families are present but more dispersed. Against this shared background, lineage-specific content is modest but asymmetric, and larger in *P. ficus*. The clearest lineage-specific signal is a *P. ficus* expansion of secreted UDP-glycosyltransferases, organized as two tandem arrays absent from *P. citri* (**Fig. 2E**). UGTs glycosylate lipophilic substrates, including plant secondary metabolites, and are a recurring component of insect xenobiotic detoxification (Ahn et al. 2012; Meech et al. 2019); tandem UGT cluster expansions have been linked to host-plant detoxification and adaptation in other insects (Ahn et al. 2012; Wang et al. 2024). The *P. ficus* expansion is notable in occurring in the narrower-host species rather than the broad-host *P. citri*, though the relationship between UGT copy number and host breadth is not consistent across insects (Wang et al. 2024).

### Preliminary assessment of genetic diversity in California *P. ficus* and markers for population monitoring

*P. ficus* is invasive in California, where prior molecular work pointed to limited genetic variation consistent with introduction from a restricted source pool (Daane et al. 2018). No published whole-genome estimates were available to our knowledge. We resequenced four California field individuals against the new reference to benchmark diversity, revisit the limited-variation inference, and identify genome-anchored SSR markers for vineyard-scale monitoring.

The genome-wide π of 2.06 x 10⁻³ is higher than would be expected from a severely depleted post-introduction gene pool. This is the first whole-genome estimate for California *P. ficus* and mildly complicates the inference of limited variation drawn from earlier marker-based work. Two interpretations are compatible with the data: the California invasion may have retained more standing variation than previously appreciated, or these four field individuals capture more diversity than is present across the broader California range. Distinguishing these possibilities will require denser sampling across vineyards and producing regions, ideally paired with samples from the Middle Eastern source range identified by Daane et al. (2018). Four individuals are a small sample, and even individuals from geographically distinct sites are likely to share substantial recent ancestry given the severity of the founder event; these estimates are therefore a first whole-genome benchmark rather than a population-level characterization. The 132 cross-validated polymorphic SSR markers are the immediate practical output (**Table S13**). Each marker is polymorphic across the four California field individuals and should be directly usable for vineyard-scale population monitoring, addressing a longstanding need for population-genetic tools in vine mealybug research (Daane et al. 2018).

### Pervasive, female-skewed sex-biased expression

Adult male and female *P. ficus* differ dramatically in morphology (**Fig. 1**) and life history. Females are large, sessile, neotenic, and feed continuously over an extended adult lifespan; males are small, winged, non-feeding, and live a few days dedicated to mate location (Daane et al. 2012; Bain et al. 2021). Mealybugs achieve this dimorphism without sex chromosomes: sex is determined by PGE, in which males are diploid at fertilization but transcriptionally silence their paternally inherited chromosomes during early development (Bain et al. 2021; de la Filia et al. 2021). We sequenced adult male and female transcriptomes to ask two questions: how is sex-biased expression organized in *P. ficus*, and how much of it reflects PGE silencing itself?

About 40% of the genes were differentially expressed between adult females and males. Female-biased loci outnumber male-biased loci 1.57-fold and show 1.7-fold larger median fold changes. A comparable female skew at the level of moderately sex-biased genes has been reported in *P. citri* (Bain et al. 2021), although in that species the most extreme bias (>10-fold) is male-directed; the same study found female- and male-biased genes both enriched for core biological processes. Male-biased genes are enriched for sensory and locomotor functions (odorant binding, nervous system, muscle, ion transport, cilium organization), and female-biased genes for core biosynthesis (translation, ribosome biogenesis, mRNA metabolism, nuclear/nucleolar components); this functional partition (male sensory, locomotor, and structural versus female biosynthetic) parallels the categories described in *M. hirsutus* (Kohli et al. 2021), although that study identified far fewer sex-biased genes and no genome-wide expression difference between the sexes. These categories are consistent with the divergent adult biology of the two sexes but do not by themselves resolve whether they reflect direct sex-specific specialization or downstream consequences of upstream regulators of dimorphism, which in mealybugs operate through PGE rather than a conventional sex-determination cascade (Bain et al. 2021; de la Filia et al. 2021).

Both sex-biased gene sets are enriched for predicted secretion relative to unbiased genes, consistent with secretion being central to the adult biology of both sexes, but the two secretomes differ in functional composition and evolutionary origin. The female-biased enrichment fits the continuous feeding of the sessile female, and the male-biased enrichment is dominated by odorant-binding, cuticle, and chitin-binding proteins, fitting the short-lived, dispersing, mate-locating male (Corcoran and Mahaffee 2024). More informative than the enrichment itself is the evolutionary origin of the two sets. The male-biased secretome is drawn preferentially from secreted proteins with 1:1 orthologs in *P. citri*, whereas the female-biased secretome spans the broader, lineage-specific effector pool and is functionally heterogeneous, with few enriched domains over a large candidate set and many members lacking any detectable InterPro signature. We read this as lineage-specific selection acting on the feeding-related secretome: the female feeding repertoire is evolving and diversifying on the *P. ficus* lineage, while the male secretome remains anchored in a conserved, shared core, consistent with the faster evolution and lower orthology of female-biased genes reported in *P. citri* (Mongue et al. 2025) and with the lineage-specific expansion of salivary effectors documented in aphids (Boulain et al. 2018).

PGE silencing offers a candidate mechanism for the female skew documented above. If paternal alleles in males were uniformly silenced without compensation, male output would be globally halved relative to diploid females, producing a genome-wide log₂FC near −1. Our data do not show this (**Fig. 3C**). The Ps median is −0.33 (mode near zero), meaning the median gene expresses about 80% of female level rather than 50%. This matches allele-specific RNA-Seq in *P. citri* and *P. citri* x *P. ficus* hybrid males, where paternal silencing in soma is strong but incomplete, with up to ∼70% of somatically expressed genes retaining some paternal contribution even as maternal alleles dominate globally (only ∼20% of genes showing any paternal contribution; de la Filia et al. 2021). Bain et al. (2021) proposed female hypermethylation in *P. citri* as a possible mechanism of ploidy compensation. Somatic paternal silencing in *P. ficus* therefore appears partial, and is not on its own sufficient to explain the female-skewed sex bias we observe.

### Genomic clustering of female-biased expression

The most striking finding from the expression analysis is that female-biased expression is not dispersed across the genome but concentrated in a small number of discrete scaffold-level blocks. The same is not true to nearly the same degree for male-biased expression. Permutation of sex-bias labels confirms that this degree of clustering is far beyond chance (**Fig. 4**, **Table 4**), so the blocks are a real feature of genome organization rather than a sampling artifact of a few gene-rich scaffolds. What makes these female blocks distinctive is not simply that they carry many female-biased genes, but that they share three properties that together point to coordinate regulation. They are unusually gene-dense; they each harbor a local expansion of a single gene family: phosphatidylethanolamine-binding proteins (PEBPs) on Ps350, glycosyl- and sulfo-transferases on Ps72, carboxylesterases on Ps743; and the female-biased genes within a block are tightly co-expressed across samples, far more so than female-biased genes drawn at random from the genome. Female-biased genes inside these blocks are also more strongly biased than female-biased genes elsewhere, and the blocks are not enriched for repetitive sequence. The blocks therefore could be interpreted as coordinately regulated genomic units rather than incidental clusters of co-located paralogs. This fits the faster evolution and lower orthology of female-biased genes reported in *P. citri* (Mongue et al. 2025) and parallels the lineage-specific tandem expansions of salivary effectors documented in aphids (Boulain et al. 2018). To our knowledge, female-restricted coordinate regulation of clustered tandem paralogs on this scale has not been described in a hemipteran lacking sex chromosomes; the chromosome-level sex-bias enrichments reported in aphids are X-linked, reflecting the distinct evolutionary dynamics of the X under X0 sex determination (Jaquiéry et al. 2013; Y. Li et al. 2020), a substrate the PGE genome of *P. ficus* does not have, since mealybug sex is determined by paternal genome elimination without sex chromosomes (Kol-Maimon et al. 2014; Bain et al. 2021).

The biological roles of these expansions remain to be studied. PEBPs on Ps350 belong to a family with established roles in MAPK/ERK signaling (Yeung et al. 1999) and account for the GO enrichment for negative regulation of the ERK1/2 cascade among female-biased genes. The Ps72 glycosyl- and sulfo-transferase cluster is plausibly involved in glycan modification of cuticular or secreted proteins. The Ps743 carboxylesterase cluster is of immediate applied interest: carboxylesterases are recurrent substrates for metabolic insecticide resistance in hemipteran pests, often via gene amplification and overexpression, as in the amplified E4/FE4 esterases of *Myzus persicae* (Bass et al. 2014) and esterase-mediated organophosphate resistance in *Nilaparvata lugens* (Small and Hemingway 2000). A female-specific, coordinately expressed esterase expansion is therefore a plausible substrate for resistance evolution under the insecticide-heavy management typical of vine mealybug (Mansour et al. 2018; Daane et al. 2020).

### Male-biased expression clusters without gene-family expansion

Male-biased expression is also clustered, but the M-enriched scaffolds differ from F-enriched ones in three respects: they are not gene-dense above the genome background, they lack obvious tandem family expansions and instead contain heterogeneous gene content (zinc finger paralogs, retrovirus-related Pol polyproteins, a CYP4g15-like P450, signaling components), and the effect sizes of their male-biased genes are statistically indistinguishable from male-biased genes elsewhere. What they share with F-blocks is strong within-block co-expression, consistent with coordinate regulation despite heterogeneous gene content. The 2,862 male-biased genes cannot be a passive consequence of uniform paternal-genome silencing. Under PGE, silencing would produce a global deficit of male transcription, not active large-effect male upregulation. These genes are actively upregulated in males despite the halved paternal-genome dose, and the M-enriched scaffolds represent regions where male-specific regulatory programs override the silencing baseline. Whether they correspond to regions of male-specific transcriptional activation, regions where paternal alleles preferentially escape silencing, or both, cannot be resolved with bulk RNA-Seq.

The asymmetry between sexes (more female-biased genes, larger female-biased effects, gene-rich tandem-expanded female blocks against heterogeneous modest-effect male blocks) fits a regulatory architecture in which the male transcriptome is more constrained than the female one. Mongue et al. (2025) reported lower variation and slower sequence evolution of male-biased genes in *P. citri*, alongside signatures of stronger adaptation; our data extend that to *P. ficus* and add a spatial dimension, in which female-biased gene evolution in mealybugs is proceeding preferentially through tandem duplication and coordinate regulation of clustered paralogs.

In summary, sex-biased expression in *P. ficus* is spatially structured, and the structure differs by sex. Female-biased genes cluster into gene-dense blocks of co-expressed tandem paralogs; male-biased genes cluster into co-expressed but compositionally heterogeneous regions without family expansion. These are descriptive genomic patterns from a single bulk RNA-Seq comparison of three females and three males; their regulatory basis, their relationship to paternal-genome elimination, and any functional consequences remain to be tested directly.

**Supplemental Material:** available at G3 online.

## Data availability

Raw sequencing reads and the genome assembly/annotations were deposited at NCBI under BioProject PRJNA1481362, PRJNA1482103 (primary scaffolds), and PRJNA1482104 (haplotigs). PASS-filtered SNP and indel VCFs together with all assembly and annotation files (fasta and gff) are available at Zenodo (10.5281/zenodo.20854017; link open to reviewers: https://zenodo.org/records/21039194?token=eyJhbGciOiJIUzUxMiJ9.eyJpZCI6ImNmZTE4MTU0LWNhMGUtNGU0ZS05NTBmLThlNTg1MTUwZDI1MSIsImRhdGEiOnt9LCJyYW5kb20iOiJhMzU0YjA2ODU1NzQ2YWFiMmI1M2JmNjM2MTIyZGU0OSJ9.PQrDj1ZIpPZeG0R_Lyq0vs-L5JQmH5y_nxuD_VxIQYpMwhDxMcFJPvf-XaQR9JuBa07xlA8CwOxv6vdJOWvYew). A dedicated genome browser is available at www.grapegenomics.com. Analysis code is available at https://github.com/CantuLab/VMB_genome_analyses.

## Author contributions

Conceptualization: D.C., L.B., R.N., M.S.

Formal analysis: D.C., A.M.

Sample processing and library preparation: R.F.-B.

Resources: M.S., N.C.

Writing-original draft preparation: D.C.

Visualization: D.C.

Project administration: D.C., L.B.

Funding acquisition: D.C., L.B., R.N., M.S.

All authors read and agreed to the published version of the manuscript.

## Acknowledgments

We thank the UC Davis DNA Technologies Core for sequencing assistance.

## References

Ahn S-J, Vogel H, Heckel DG. 2012. Comparative analysis of the UDP-glycosyltransferase multigene family in insects. Insect Biochem Mol Biol. 42(2):133–147. 10.1016/j.ibmb.2011.11.006

Almeida RPP et al. 2013. Ecology and management of grapevine leafroll disease. Front Microbiol. 4:94. 10.3389/fmicb.2013.00094

Bain SA et al. 2021. Sex-specific expression and DNA methylation in a species with extreme sexual dimorphism and paternal genome elimination. Mol Ecol. 30(22):5687–5703. 10.1111/mec.15842

Bankevich A et al. 2012. SPAdes: A New Genome Assembly Algorithm and Its Applications to Single-Cell Sequencing. J Comput Biol. 19(5):455–477. 10.1089/cmb.2012.0021

Bass C et al. 2014. The evolution of insecticide resistance in the peach potato aphid, *Myzus persicae*. Insect Biochem Mol Biol. 51:41–51. 10.1016/j.ibmb.2014.05.003

Beier S et al. 2017. MISA-web: a web server for microsatellite prediction. Bioinformatics. 33(16):2583–2585. 10.1093/bioinformatics/btx198

Boetzer M, Pirovano W. 2014. SSPACE-LongRead: scaffolding bacterial draft genomes using long read sequence information. BMC Bioinformatics. 15(1):211. 10.1186/1471-2105-15-211

Bolger AM, Lohse M, Usadel B. 2014. Trimmomatic: a flexible trimmer for Illumina sequence data. Bioinformatics. 30(15):2114–2120. 10.1093/bioinformatics/btu170

Bongiorni S, Fiorenzo P, Pippoletti D, Prantera G. 2004. Inverted meiosis and meiotic drive in mealybugs. Chromosoma. 112(7):331–341. 10.1007/s00412-004-0278-4

Boratyn GM et al. 2019. Magic-BLAST, an accurate RNA-seq aligner for long and short reads. BMC Bioinformatics. 20(1):405. 10.1186/s12859-019-2996-x

Borowiec ML. 2016. AMAS: a fast tool for alignment manipulation and computing of summary statistics. PeerJ. 4:e1660. 10.7717/peerj.1660

Boulain H et al. 2018. Fast Evolution and Lineage-Specific Gene Family Expansions of Aphid Salivary Effectors Driven by Interactions with Host-Plants. Genome Biol Evol. 10(6):1554–1572. 10.1093/gbe/evy097

Brun LO et al. 1995. Functional haplodiploidy: a mechanism for the spread of insecticide resistance in an important international insect pest. Proc Natl Acad Sci. 92(21):9861–9865. 10.1073/pnas.92.21.9861

Buchfink B, Reuter K, Drost H-G. 2021. Sensitive protein alignments at tree-of-life scale using DIAMOND. Nat Methods. 18(4):366–368. 10.1038/s41592-021-01101-x

Camacho C et al. 2009. BLAST+: architecture and applications. BMC Bioinformatics. 10(1):421. 10.1186/1471-2105-10-421

Capella-Gutiérrez S, Silla-Martínez JM, Gabaldón T. 2009. trimAl: a tool for automated alignment trimming in large-scale phylogenetic analyses. Bioinformatics. 25(15):1972–1973. 10.1093/bioinformatics/btp348

Carolan JC et al. 2011. Predicted Effector Molecules in the Salivary Secretome of the Pea Aphid (Acyrthosiphon pisum): A Dual Transcriptomic/Proteomic Approach. J Proteome Res. 10(4):1505–1518. 10.1021/pr100881q

Carrière Y. 2003. Haplodiploidy, Sex, and the Evolution of Pesticide Resistance. J Econ Entomol. 96(6):1626–1640. 10.1093/jee/96.6.1626

Chin C-S et al. 2016. Phased diploid genome assembly with single-molecule real-time sequencing. Nat Methods. 13(12):1050–1054. 10.1038/nmeth.4035

Choi J, Lee S. 2022. Higher classification of mealybugs (Hemiptera: Coccomorpha) inferred from molecular phylogeny and their endosymbionts. Syst Entomol. 47(2):354–370. 10.1111/syen.12534

Cocco A et al. 2021. Sustainable management of the vine mealybug in organic vineyards. J Pest Sci. 94(2):153–185. 10.1007/s10340-020-01305-8

Cochetel N et al. 2021. Diploid chromosome-scale assembly of the Muscadinia rotundifolia genome supports chromosome fusion and disease resistance gene expansion during Vitis and Muscadinia divergence. G3 GenesGenomesGenetics. 11(4):jkab033. 10.1093/g3journal/jkab033

Corcoran JA, Mahaffee WF. 2024. Identification of a receptor for the sex pheromone of the vine mealybug, Planococcus ficus. Curr Res Insect Sci. 5:100072. 10.1016/j.cris.2024.100072

Daane KM et al. 2011. Development of a Multiplex Pcr for Identification of Vineyard Mealybugs. Environ Entomol. 40(6):1595–1603. 10.1603/EN11075

Daane KM et al. 2012. Biology and Management of Mealybugs in Vineyards. In: Bostanian NJ, Vincent C, Isaacs R, editors. Arthropod Management in Vineyards: Pests, Approaches, and Future Directions. Springer Netherlands; p 271–307 [accessed 2026 June 11]. 10.1007/978-94-007-4032-7_12.

Daane KM et al. 2018. Determining the geographic origin of invasive populations of the mealybug Planococcus ficus based on molecular genetic analysis. PLOS ONE. 13(3):e0193852. 10.1371/journal.pone.0193852

Daane KM et al. 2020. Development of a Mating Disruption Program for a Mealybug, Planococcus ficus, in Vineyards. Insects. 11(9):635. 10.3390/insects11090635

Danecek P et al. 2021. Twelve years of SAMtools and BCFtools. GigaScience. 10(2):giab008. 10.1093/gigascience/giab008

Deng J et al. 2025. Genomic insights into the phylogeny and evolutionary history of scale insects (Hemiptera: Coccoidea): Resolving family-level relationships. Mol Phylogenet Evol. 210:108383. 10.1016/j.ympev.2025.108383

Elzinga DA, Jander G. 2013. The role of protein effectors in plant–aphid interactions. Curr Opin Plant Biol. 16(4):451–456. 10.1016/j.pbi.2013.06.018

de la Filia AG et al. 2021. Males That Silence Their Father’s Genes: Genomic Imprinting of a Complete Haploid Genome. Mol Biol Evol. 38(6):2566–2581. 10.1093/molbev/msab052

de la Filia AG, Bain SA, Ross L. 2015. Haplodiploidy and the reproductive ecology of Arthropods. Curr Opin Insect Sci. 9:36–43 (Pests and resistance * Behavioural ecology). 10.1016/j.cois.2015.04.018

de la Filia AG, Fenn-Moltu G, Ross L. 2019. No evidence for an intragenomic arms race under paternal genome elimination in Planococcus mealybugs. J Evol Biol. 32(5):491–504. 10.1111/jeb.13431

Garcia-Morales M et al. 2016. ScaleNet: a literature-based model of scale insect biology and systematics. Database. [published online ahead of print] [accessed 2026 June 3]. https://academic.oup.com/database/article/doi/10.1093/database/bav118/2630093/ScaleNet-a-literaturebased-model-of-scale-insect. 10.1093/database/bav118

Gardner A, Ross L. 2014. Mating ecology explains patterns of genome elimination. Ecol Lett. 17(12):1602–1612. 10.1111/ele.12383

Gotz S et al. 2008. High-throughput functional annotation and data mining with the Blast2GO suite. Nucleic Acids Res. 36(10):3420–3435. 10.1093/nar/gkn176

Haas BJ et al. 2003. Improving the Arabidopsis genome annotation using maximal transcript alignment assemblies. Nucleic Acids Res. 31(19):5654–5666. 10.1093/nar/gkg770

Haas BJ et al. 2008. Automated eukaryotic gene structure annotation using EVidenceModeler and the Program to Assemble Spliced Alignments. Genome Biol. 9(1):R7. 10.1186/gb-2008-9-1-r7

Haas BJ et al. 2013. De novo transcript sequence reconstruction from RNA-seq using the Trinity platform for reference generation and analysis. Nat Protoc. 8(8):1494–1512. 10.1038/nprot.2013.084

Hardy NB, Gullan PJ, Hodgson CJ. 2008. A subfamily-level classification of mealybugs (Hemiptera: Pseudococcidae) based on integrated molecular and morphological data. Syst Entomol. 33(1):51–71. 10.1111/j.1365-3113.2007.00408.x

Herbette M, Ross L. 2023. Paternal genome elimination: patterns and mechanisms of drive and silencing. Curr Opin Genet Dev. 81:102065. 10.1016/j.gde.2023.102065

Jaquiéry J et al. 2013. Masculinization of the X Chromosome in the Pea Aphid. PLOS Genet. 9(8):e1003690. 10.1371/journal.pgen.1003690

Jones P et al. 2014. InterProScan 5: genome-scale protein function classification. Bioinformatics. 30(9):1236–1240. 10.1093/bioinformatics/btu031

Kent WJ. 2002. BLAT—The BLAST-Like Alignment Tool. Genome Res. 12(4):656–664. 10.1101/gr.229202

Kim D et al. 2019. Graph-based genome alignment and genotyping with HISAT2 and HISAT-genotype. Nat Biotechnol. 37(8):907–915. 10.1038/s41587-019-0201-4

Kim D, Langmead B, Salzberg SL. 2015. HISAT: a fast spliced aligner with low memory requirements. Nat Methods. 12(4):357–360. 10.1038/nmeth.3317

Kohli S et al. 2021. Genome and transcriptome analysis of the mealybug *Maconellicoccus hirsutus*: Correlation with its unique phenotypes. Genomics. 113(4):2483–2494. 10.1016/j.ygeno.2021.05.014

Kol-Maimon H, Mendel Z, Franco JC, Ghanim M. 2014. Paternal inheritance in mealybugs (Hemiptera: Coccoidea: Pseudococcidae). Naturwissenschaften. 101(10):791–802. 10.1007/s00114-014-1218-7

Korf I. 2004. Gene finding in novel genomes. BMC Bioinformatics. 5:59. 10.1186/1471-2105-5-59

Li H. 2013. Aligning sequence reads, clone sequences and assembly contigs with BWA-MEM. [accessed 2026 June 15]. http://arxiv.org/abs/1303.3997. 10.48550/arXiv.1303.3997

Li H. 2021. New strategies to improve minimap2 alignment accuracy. Bioinformatics. 37(23):4572–4574. 10.1093/bioinformatics/btab705

Li H, Durbin R. 2009. Fast and accurate short read alignment with Burrows–Wheeler transform. Bioinformatics. 25(14):1754–1760. 10.1093/bioinformatics/btp324

Li M et al. 2020. A chromosome-level genome assembly provides new insights into paternal genome elimination in the cotton mealybug Phenacoccus solenopsis. Mol Ecol Resour. 20(6):1733–1747. 10.1111/1755-0998.13232

Li Y, Zhang B, Moran NA. 2020. The Aphid X Chromosome Is a Dangerous Place for Functionally Important Genes: Diverse Evolution of Hemipteran Genomes Based on Chromosome-Level Assemblies. Mol Biol Evol. 37(8):2357–2368. 10.1093/molbev/msaa095

Lomsadze A, Ter-Hovhannisyan V, Chernoff YO, Borodovsky M. 2005. Gene identification in novel eukaryotic genomes by self-training algorithm. Nucleic Acids Res. 33(20):6494–6506. 10.1093/nar/gki937

Love MI, Huber W, Anders S. 2014. Moderated estimation of fold change and dispersion for RNA-seq data with DESeq2. Genome Biol. 15(12):550. 10.1186/s13059-014-0550-8

Ma R, Reese JC, Black WC, Bramel-Cox P. 1990. Detection of pectinesterase and polygalacturonase from salivary secretions of living greenbugs, *Schizaphis graminum* (Homoptera: Aphididae). J Insect Physiol. 36(7):507–512. 10.1016/0022-1910(90)90102-L

Manni M et al. 2021. BUSCO Update: Novel and Streamlined Workflows along with Broader and Deeper Phylogenetic Coverage for Scoring of Eukaryotic, Prokaryotic, and Viral Genomes. Mol Biol Evol. 38(10):4647–4654. 10.1093/molbev/msab199

Mansour R et al. 2018. Vine and citrus mealybug pest control based on synthetic chemicals. A review. Agron Sustain Dev. 38(4):37. 10.1007/s13593-018-0513-7

Marçais G, Kingsford C. 2011. A fast, lock-free approach for efficient parallel counting of occurrences of k-mers. Bioinformatics. 27(6):764–770. 10.1093/bioinformatics/btr011

McAllan JW, Adams JB. 1961. The significance of pectinase in plant penetration by aphids. Can J Zool. 39(3):305–310. 10.1139/z61-034

McKenna A et al. 2010. The Genome Analysis Toolkit: a MapReduce framework for analyzing next-generation DNA sequencing data. Genome Res. 20(9):1297–1303. 10.1101/gr.107524.110

Meech R et al. 2019. The UDP-Glycosyltransferase (UGT) Superfamily: New Members, New Functions, and Novel Paradigms. Physiol Rev. 99(2):1153–1222. 10.1152/physrev.00058.2017

Miles A et al. 2024. cggh/scikit-allel: v1.3.13. [accessed 2026 June 15]. https://zenodo.org/records/13772087. 10.5281/zenodo.13772087

Minh BQ et al. 2020. IQ-TREE 2: New Models and Efficient Methods for Phylogenetic Inference in the Genomic Era. Mol Biol Evol. 37(5):1530–1534. 10.1093/molbev/msaa015

Misof B et al. 2014. Phylogenomics resolves the timing and pattern of insect evolution. Science. 346(6210):763–767. 10.1126/science.1257570

Mongue AJ et al. 2025. Contrasting Evolutionary Trajectories Under Paternal Genome Elimination in Male and Female Citrus Mealybugs. Mol Ecol. 34(13):e17826. 10.1111/mec.17826

Mose L. 2025. mozack/ubu. [accessed 2026 June 15]. https://github.com/mozack/ubu

Patro R et al. 2017. Salmon provides fast and bias-aware quantification of transcript expression. Nat Methods. 14(4):417–419. 10.1038/nmeth.4197

Pertea M et al. 2015. StringTie enables improved reconstruction of a transcriptome from RNA-seq reads. Nat Biotechnol. 33(3):290–295. 10.1038/nbt.3122

Ranallo-Benavidez TR, Jaron KS, Schatz MC. 2020. GenomeScope 2.0 and Smudgeplot for reference-free profiling of polyploid genomes. Nat Commun. 11(1):1432. 10.1038/s41467-020-14998-3

dos Reis M, Yang Z. 2011. Approximate likelihood calculation on a phylogeny for Bayesian estimation of divergence times. Mol Biol Evol. 28(7):2161–2172. 10.1093/molbev/msr045

Ross L, Mongue AJ, De La Filia A. 2024. The genome sequence of the citrus … Wellcome Open Res. [published online ahead of print] [accessed 2026 May 30]. https://wellcomeopenresearch.org/articles/9-22

Sharma A et al. 2014. Salivary proteins of plant-feeding hemipteroids – implication in phytophagy. Bull Entomol Res. 104(2):117–136. 10.1017/S0007485313000618

Shi G et al. 2012. Age constraint on Burmese amber based on U–Pb dating of zircons. Cretac Res. 37:155–163. 10.1016/j.cretres.2012.03.014

Silva-Sanzana C, Estevez JM, Blanco-Herrera F. 2020. Influence of cell wall polymers and their modifying enzymes during plant–aphid interactions. J Exp Bot. 71(13):3854–3864. 10.1093/jxb/erz550

Sisterson MS, Uchima SY. 2024. Planococcus ficus (Hemiptera: Pseudococcidae) movement and demography: methods for generating cohorts for laboratory studies. J Econ Entomol. 117(1):118–126. 10.1093/jee/toad210

Slater GSC, Birney E. 2005. Automated generation of heuristics for biological sequence comparison. BMC Bioinformatics. 6:31. 10.1186/1471-2105-6-31

Small GJ, Hemingway J. 2000. Molecular characterization of the amplified carboxylesterase gene associated with organophosphorus insecticide resistance in the brown planthopper, Nilaparvata lugens. Insect Mol Biol. 9(6):647–653. 10.1046/j.1365-2583.2000.00229.x

Soneson C, Love MI, Robinson MD. 2016. Differential analyses for RNA-seq: transcript-level estimates improve gene-level inferences. F1000Research. 4:1521. 10.12688/f1000research.7563.2

Stanke M et al. 2006. AUGUSTUS: ab initio prediction of alternative transcripts. Nucleic Acids Res. 34(Web Server issue):W435–439. 10.1093/nar/gkl200

Teufel F et al. 2022. SignalP 6.0 predicts all five types of signal peptides using protein language models. Nat Biotechnol. 40(7):1023–1025. 10.1038/s41587-021-01156-3

TransDecoder. GitHub; [accessed 2026 June 3]. https://github.com/TransDecoder

Tsai C-W et al. 2008. Transmission of grapevine leafroll-associated virus 3 by the vine mealybug (Planococcus ficus). Phytopathology. 98(10):1093–1098. 10.1094/PHYTO-98-10-1093

Tsai C-W et al. 2010. Mealybug transmission of Grapevine leafroll viruses: an analysis of virus-vector specificity. Phytopathology. 100(8):830–834. 10.1094/PHYTO-100-8-0830

Untergasser A et al. 2012. Primer3—new capabilities and interfaces. Nucleic Acids Res. 40(15):e115. 10.1093/nar/gks596

Van Dongen S. 2008. Graph Clustering Via a Discrete Uncoupling Process. SIAM J Matrix Anal Appl. 30(1):121–141. 10.1137/040608635

Vea I, Grimaldi D. 2015. Diverse New Scale Insects (Hemiptera: Coccoidea) in Amber from the Cretaceous and Eocene with a Phylogenetic Framework for Fossil Coccoidea. Am Mus Novit. 3823. 10.1206/3823.1

Vea IM, Grimaldi DA. 2016. Putting scales into evolutionary time: the divergence of major scale insect lineages (Hemiptera) predates the radiation of modern angiosperm hosts. Sci Rep. 6(1):23487. 10.1038/srep23487

Wang H et al. 2024. UDP-glycosyltransferases act as key determinants of host plant range in generalist and specialist Spodoptera species. Proc Natl Acad Sci. 121(19):e2402045121. 10.1073/pnas.2402045121

Wang Y et al. 2012. MCScanX: a toolkit for detection and evolutionary analysis of gene synteny and collinearity. Nucleic Acids Res. 40(7):e49. 10.1093/nar/gkr1293

Will T, Furch AC, Zimmermann MR. 2013. How phloem-feeding insects face the challenge of phloem-located defenses. Front Plant Sci. 4 [accessed 2026 May 30]. https://www.frontiersin.org/journals/plant-science/articles/10.3389/fpls.2013.00336/full. 10.3389/fpls.2013.00336

Will T, Tjallingii WF, Thönnessen A, van Bel AJE. 2007. Molecular sabotage of plant defense by aphid saliva. Proc Natl Acad Sci. 104(25):10536–10541. 10.1073/pnas.0703535104

Wu TD, Watanabe CK. 2005. GMAP: a genomic mapping and alignment program for mRNA and EST sequences. Bioinformatics. 21(9):1859–1875. 10.1093/bioinformatics/bti310

Yang Z. 2007. PAML 4: phylogenetic analysis by maximum likelihood. Mol Biol Evol. 24(8):1586–1591. 10.1093/molbev/msm088

Yeung K et al. 1999. Suppression of Raf-1 kinase activity and MAP kinase signalling by RKIP. Nature. 401(6749):173–177. 10.1038/43686

Zhang P-J et al. 2015. The mealybug Phenacoccus solenopsis suppresses plant defense responses by manipulating JA-SA crosstalk. Sci Rep. 5(1):9354. 10.1038/srep09354

