## Supplementary Figures for "Genomic compartmentalization of pervasive sex-biased gene expression in the vine mealybug *Planococcus ficus*"

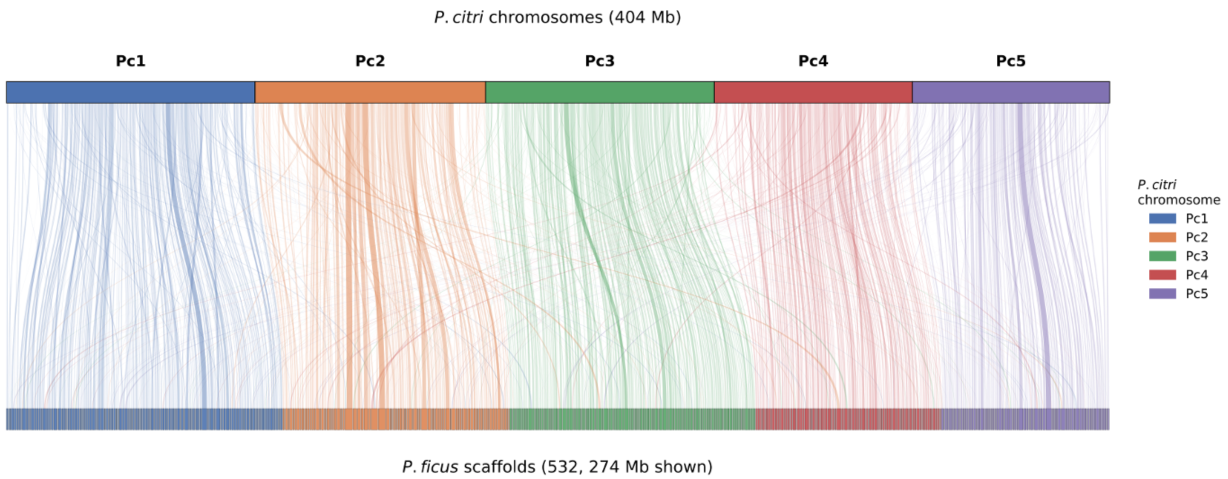


**Figure S1. Macrosynteny between *Planococcus ficus* and *P. citri*.** Top bar: the five chromosomes of the *P. citri* reference assembly (GCF_950023065.1; 403.6 Mb total). Bottom bar: 532 *P. ficus* scaffolds with ≥100 kb aligned (in alignments ≥2 kb) to a single *P. citri* chromosome, ordered by their primary syntenic correspondence and colored by that assignment. Ribbons connect chained minimap2 syntenic blocks of ≥30 kb; ribbon opacity scales with block length.

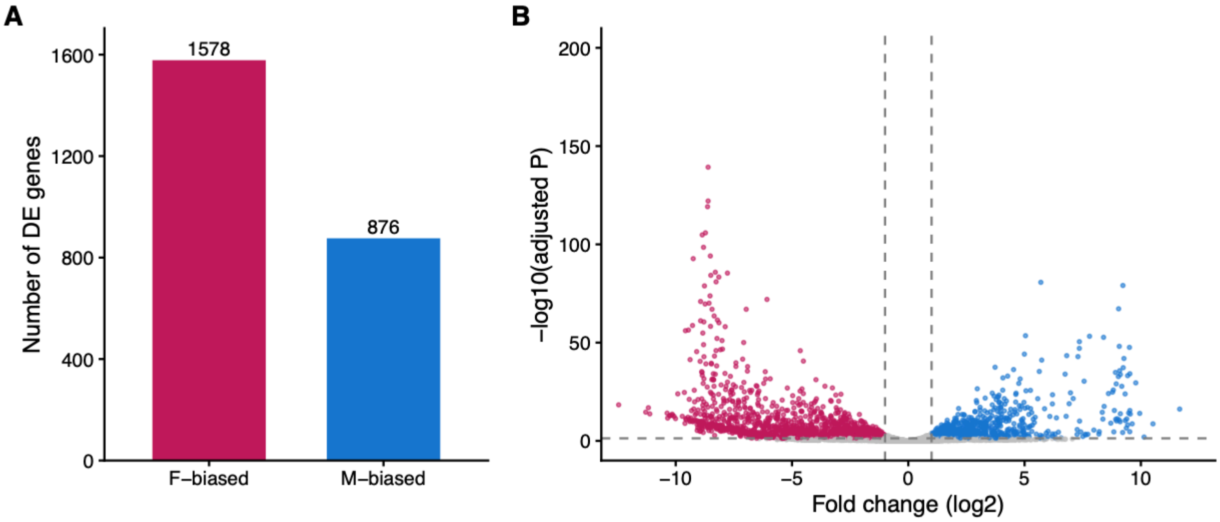


**Figure S2. Sex-biased differential expression on the haplotig (Hc) partition of the Planococcus ficus genome.** Differential expression between adult females and males for genes annotated on retained haplotigs (Hc). **(A)** Number of female-biased and male-biased Hc genes ($|\log_2 FC| > 1$, BH-adjusted $P < 0.05$; 1,578 F-biased, 876 M-biased). **(B)** Volcano plot of $\log_2$ fold change (male/female) against −log10 BH-adjusted P for all DE-tested Hc genes.

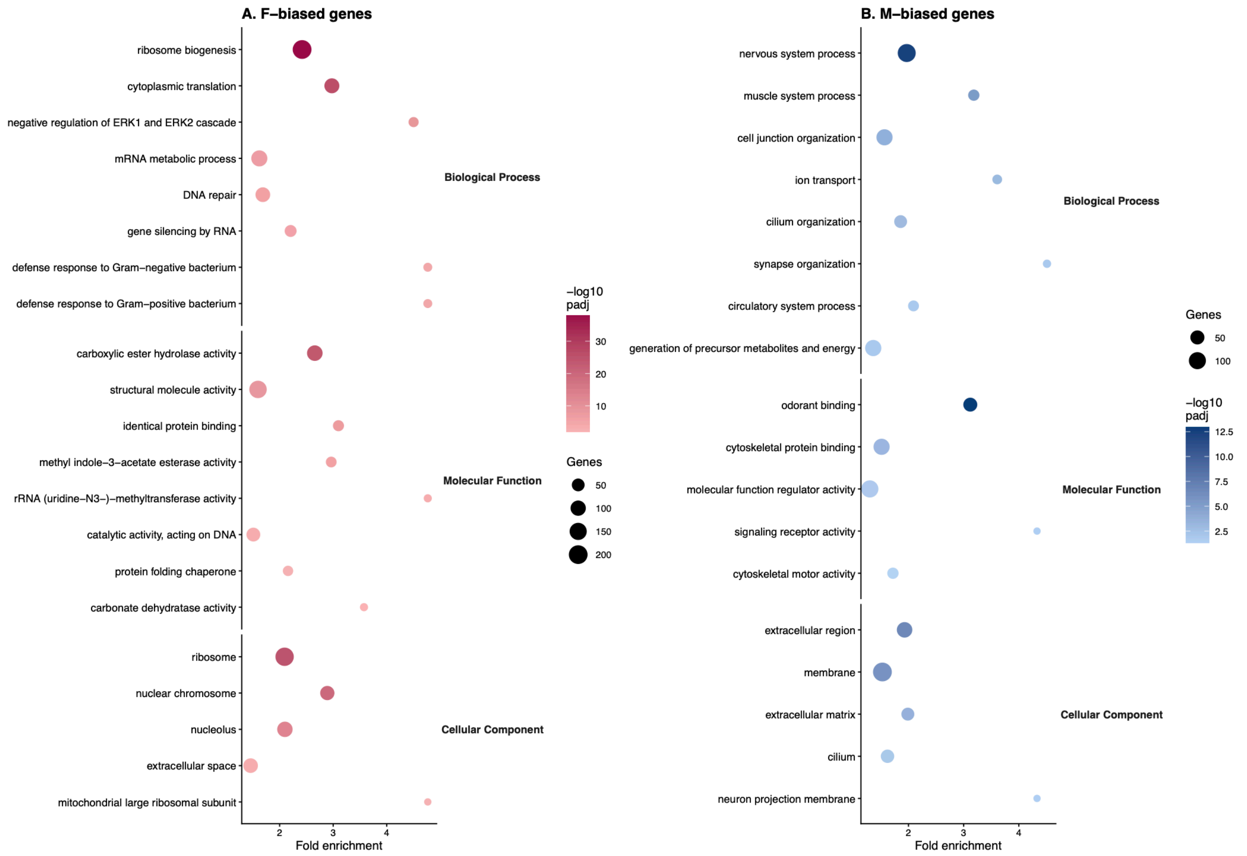


**Figure S3. Gene Ontology enrichment among sex-biased Planococcus ficus genes.** GO terms over-represented (one-sided Fisher's exact test, BH-adjusted $P < 0.05$) among (A) female-biased and (B) male-biased genes, grouped by ontology (Biological Process, Molecular Function, Cellular Component). Full values in **Table S15**.

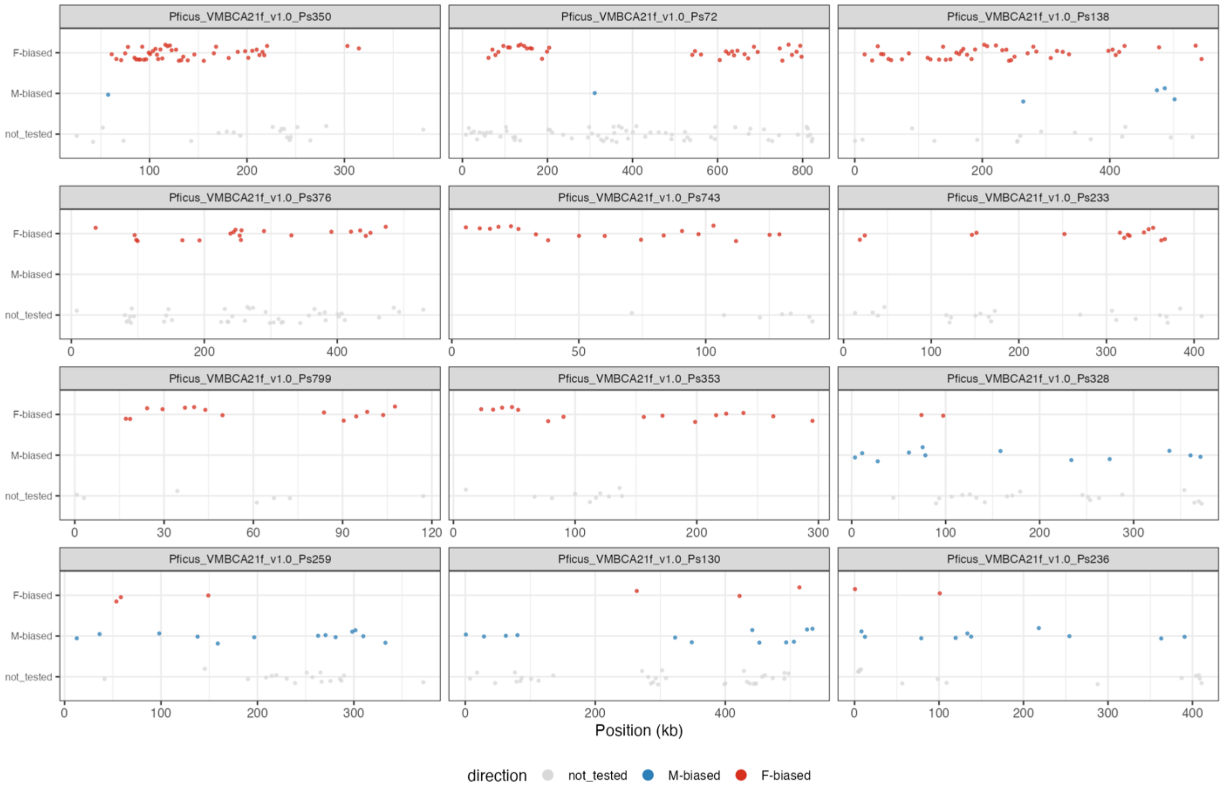


**Figure S4. Physical clustering of sex-biased genes on enriched scaffolds.** Each panel plots the midpoint position (in kb) of every gene on one enriched scaffold. Twelve scaffolds are shown: eight enriched for female-biased expression and four for male-biased expression (six from the two-sided main test, **Table 4**, and six from the directional one-sided test, **Table S16**). Genes are coloured by class: F-biased (red), M-biased (blue), or not classified as sex-biased (not_tested, grey).
